# The genes controlling normal function of citrate and spermine secretion is lost in aggressive prostate cancer and prostate model systems

**DOI:** 10.1101/2021.09.21.461176

**Authors:** Morten Beck Rye, Sebastian Krossa, Martina Hall, Casper van Mourik, Tone F. Bathen, Finn Drabløs, May-Britt Tessem, Helena Bertilsson

## Abstract

**Background:** Secretion of the metabolites citrate and spermine into prostate lumen is a unique hallmark for normal prostate epithelial cells. However, the identity of the genes controlling citrate and spermine secretion remains mostly unknown despite their obvious relevance for progression to aggressive prostate cancer.

**Materials & Methods:** In this study, we have correlated simultaneous measurement of citrate/spermine and transcriptomics data. We have refined these gene correlations in 12 prostate cancer cohorts containing 2915 tissue samples to create a novel gene signature of 150 genes connected with citrate and spermine secretion. We further explored the signature in public data, interrogating over 18 000 samples from various tissues and model systems, including 3826 samples from prostate and prostate cancer.

**Results:** In prostate cancer, the expression of this gene signature is gradually lost in tissue from normal epithelial cells through PIN, low grade (Gleason <= 7), high grade cancer (Gleason >= 8) and metastatic lesions. The accuracy of the signature is validated by its unique enrichment in prostate compared to other tissues, and its strong enrichment in epithelial tissue compartments compared to stroma. Several zinc-binding proteins that are not previously investigated in the prostate are present in the gene signature, suggesting new mechanisms for controlling zinc homeostasis in citrate/spermine secretion. However, the absence of the gene signature in all common prostate normal and cancer cell-lines, as well as prostate organoids, underlines the challenge to study the role of these genes during prostate cancer progression in model systems.

**Conclusions:** A large collection of transcriptomics data integrated with metabolomics identifies the genes related to citrate and spermine secretion in the prostate, and show that the expression of these genes gradually decreases on the path towards aggressive prostate cancer. In addition, the study questions the relevance of currently available model systems to study metabolism in prostate cancer development.

## Introduction

A unique hallmark for normal prostate epithelial cells is their ability to secrete large amounts of the metabolites citrate. While cells from other human organs convert citrate to isocitrate for oxidation and energy production in the TCA cycle, citrate conversion in the prostate epithelium is partly blocked, leading to accumulation of citrate [1]. It has been reported that citrate/isocitrate ratios in the prostate are 40/1, compared to 10/1 in other normal organs [2]. In the prostate, excess citrate is secreted out of the cell into the prostate lumen of the prostatic glands, where it is a major constituent of the prostatic fluid and essential for normal prostate organ function.

Almost all types of prostate cancer originates from prostate epithelial cells, and nearly all prostate cancer cells lose their ability to secrete citrate during the development of malignant phenotypes [1]. A reduced level of citrate has therefore been suggested as a biomarker, both for cancer detection and for identification of cancers with more aggressive phenotypes [3-5]. Even with these demonstrations of importance for normal prostate function and prostate cancer progression, the research in this field has been markedly neglected, and thus the molecular mechanisms controlling these functions remain mostly unknown. Nevertheless, a few dedicated research groups have managed to gain some insights: First, high mitochondrial levels of zinc in the prostate block citrate conversion, and the regulation of zinc transporter genes, in particular ZIP1 (*SLC39A1*), have been suggested to play a role in regulating prostate zinc levels [6, 7]. Second, the metabolite aspartate is a possible source of increased citrate in prostate cells [8, 9]. Aspartate is a precursor for oxaloacetate, which again is a precursor of citrate synthesis, and works as a carrier of the carbon groups oxidized stepwise in the mitochondria during the TCA cycle. Increased levels of aspartate could thus lead to increased levels of citrate when the citrate/isocitrate conversion is blocked. Third, metabolites synthesized in the polyamine pathway are regulated differently in the prostate [10-12]. Some interest has been directed to the polyamine spermine, due to its extremely high correlation with citrate [13]. Spermine and citrate have opposite polarity, and it has been speculated as to whether they may form a complex which is secreted into the prostate lumen [13, 14].

These presented mechanisms, though intriguing, represent only small steps on the way to a widened perspective of this unique prostate function which is not completely understood. This includes the genes and proteins involved, as well as the connections between them. Most mechanisms suggested so far have been studied in cell-lines and animal models, which are not able to fully capture the complex interplay within human prostate tissue. However, a large amount of genome-wide –omics data on prostate tissue, as well as their model systems, has accumulated since the introduction of the genomic era more than 20 years ago. These data resources have yet to be fully exploited for research on citrate and spermine secretion in the prostate. In this study we address this hallmark property of the prostate by performing extensive bioinformatic analysis on data resources from the public domain. We do this by first creating and a robust gene signature representing citrate and spermine secretion in the prostate. We then explore and validate this gene signature in multiple datasets, cohorts and public resources. In total we analyzed 32 datasets [15-42] with 18020 samples, of which 3826 were prostate samples including normal/normal-adjacent (epithelium and stroma) tissue, high/low-grade cancer tissue, metastatic samples and model systems. Our study identifies genes strongly associated with citrate and spermine secretion in the prostate and explore their behavior during cancer progression and in model systems. Our results validate previously suggested mechanisms, but also discover new central genes and mechanisms related to citrate and spermine with potential importance for this unique and intriguing prostate hallmark.

## Results

All datasets used in this study are listed in Supplementary File 1. It includes dataset ID and abbreviation used in the main text, sample description, dataset accession and references.

### Metabolite concentrations between citrate and spermine are highly correlated across tissue samples

Our research group has previously generated a dataset with 129 normal and cancer tissue samples from the prostate, where concentrations from 23 metabolites and the expression of 14161 unique genes were measured on the exact same samples (*Bertilsson*, dataset ID 1). Similar to a previous report [12], we observed a strong correlation (r=0.95) between concentrations of the metabolites citrate and spermine across the samples (Figure 1A and B). To simplify calculations, we will use the average concentration profile for citrate and spermine throughout the rest of this study. (Figure 1C). We use the abbreviation CS (Citrate and Spermine) to denote all our results based on this average concentration profile.

**Figure 1:**
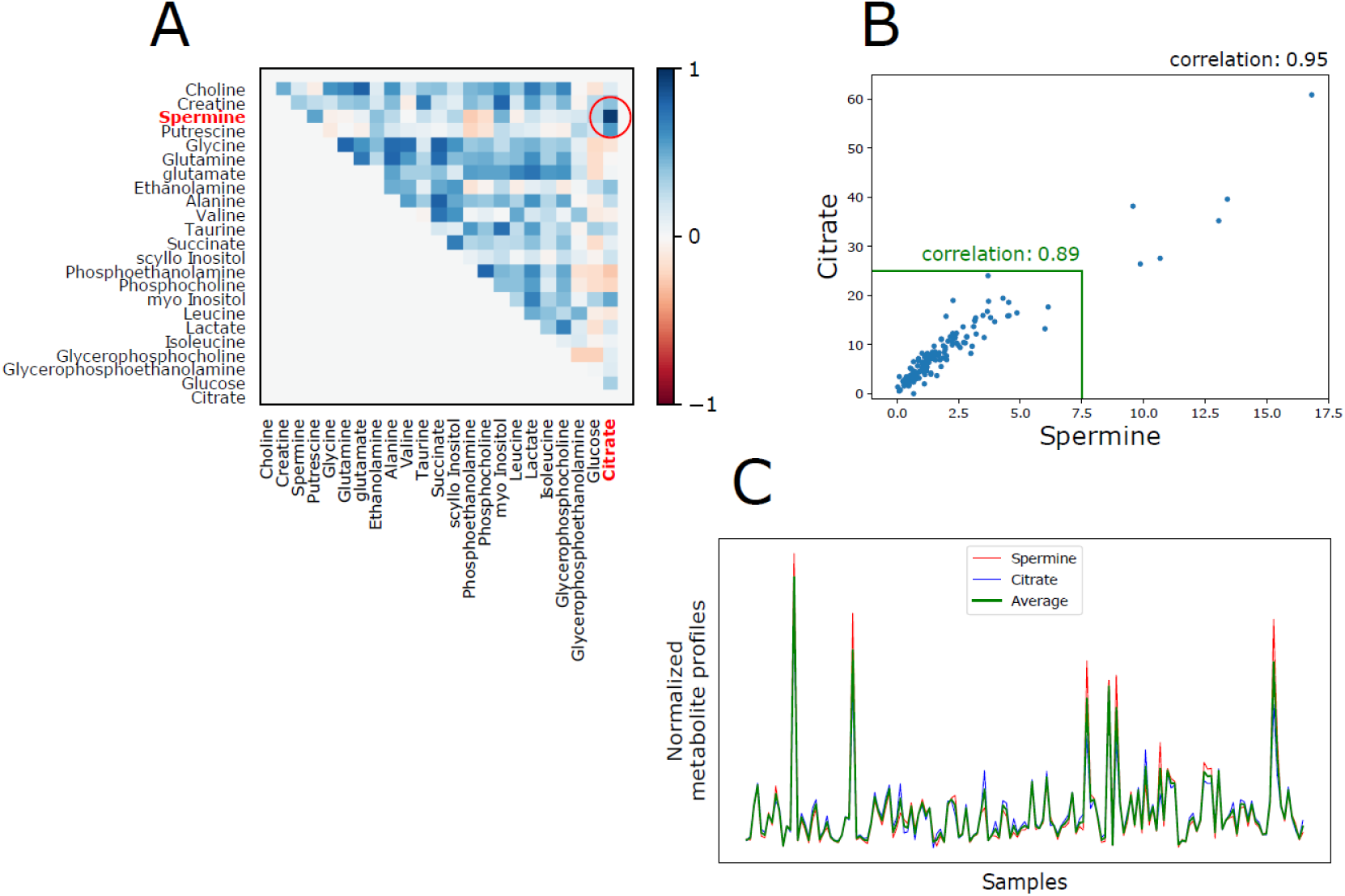
Correlations between metabolites citrate and spermine. A) Correlations between all 23 metabolite measurements across 129 prostate normal and cancer samples from the Bertilsson dataset (Dataset ID 1). The correlation is particularly strong for metabolites citrate and spermine. B) Correlations between citrate and spermine across 129 prostate normal and cancer samples from Bertilsson. The correlation is high also when extreme values are excluded (highlighted inside the green boundary). C) Similarity between the citrate, spermine and the average citrate/spermine metabolite profiles across 129 prostate normal and cancer samples from Bertilsson.

### A gene signature associated with the CS concentrations is validated in 12 datasets of prostate cancer and normal samples

We identified genes associated with the average CS levels in the prostate by calculating the Pearson correlations between the CS concentration profile and each of the 14161 unique genes measured in *Bertilsson* (dataset ID 1). The correlation analyses were performed on the 95 cancer samples in the dataset. We selected the 150 genes with the highest positive correlation which we termed the initial gene module. We further assessed the integrity of this gene module by creating a Correlation Module Score (CMS, see methods) where a high CMS means that strong intra-correlation is present between the genes in the module, resulting in a high module integrity. On the contrary, a low CMS indicates that there is low correlation between the genes in the module, and that the module integrity is lost. Moreover, if the gene module has a significantly high CMS in a dataset, it indicates that the biological process(es) which this module represents is functionally active for the samples in this dataset.

To test for significance, we generated 100 random CMSs by shuffling the CS metabolite levels between the respective samples in *Bertilsson* (dataset ID 1). The resulting CMS was statistically significant when compared to CMSs from random gene modules (CMS=0.34, p=0.007, lognormal distribution) (Figure 2A).

**Figure 2:**
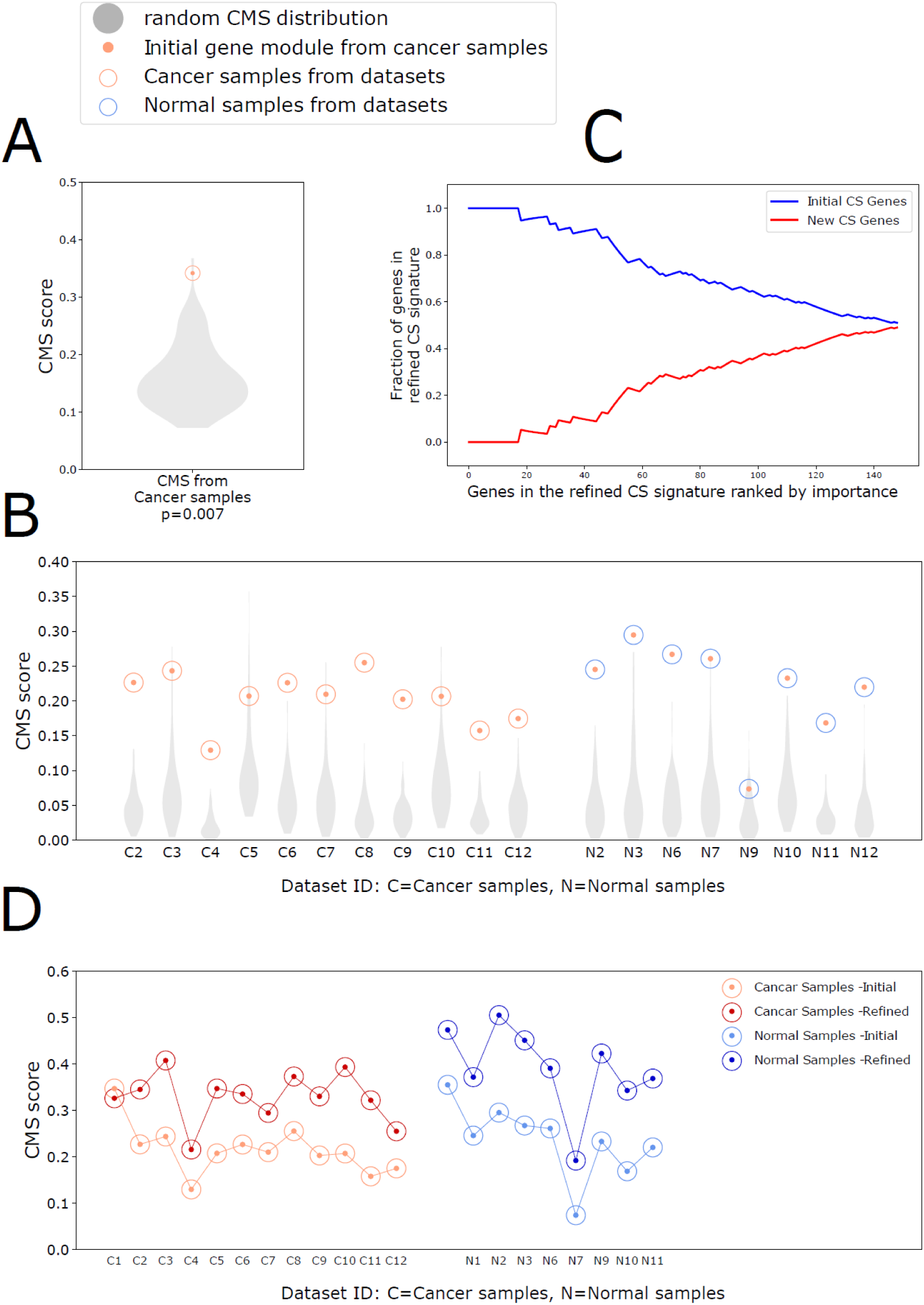
Integrity of initial and refined citrate – spermine (CS) gene signatures. A) CMS (Correlation Module Scores) for the 150 genes most positively correlated with the CS metabolite profile for cancer samples in the Bertilsson dataset (Dataset ID 1). The CMS for was statistically significant when compared to CMSs using random metabolite profiles (lognorm test) B) CMS for the initial cancer CS gene module from Bertilsson (red dots) evaluated in prostate cancer (red circles) and normal (blue circles) samples from 11 additional datasets. The CMS was statistically significant in all datasets (lognorm test, Supplementary Table T1 and T2). C) Fraction of new genes in the refined CS gene signature that replaced genes in the initial CS gene signature. The genes is sorted by importance (rank) on the bottom axis. Of the 150 top ranked genes in the initial CS signature, 74 were replaced in the refined CS signature. D) Increase in CMS after refinement of the initial CS signature for prostate normal and cancer samples in 11 datasets. The CMS increased in the normal samples, even though the normal samples were not used to create or refine the signature. Cohort ID 4, 5 and 8 did not include normal samples.

We then validated whether the integrity of the initial gene module was significantly preserved in 11 additional datasets (dataset ID 2-12), including a total of 2638 tissue samples from normal prostate and prostate cancer (ref dataset list). We calculated and compared the CMSs for the initial module, as well as the 100 random modules in all 11 datasets. The CMS from the initial gene module was validated in all 11 cancer datasets, and 7 out of 8 datasets with normal samples (lognormal test) (Figure 2B, Supplementary Table 1 and 2). We thus defined the initial module as our initial gene signature associated with CS concentrations in the prostate. We used this initial CS signature as a starting point for further refinement.

We also performed the above procedure using the 40 normal samples in *Bertilsson*, however the resulting gene module was not significant (CMS=0.37, p=0.14, lognormal distribution), and could not be validated in any of the additional datasets (Supplementary Figure 1, Supplementary Table 3 and 4). The reason for this is probably due to an insufficient number of normal samples in *Bertilsson* to perform a robust correlation analysis.

### Refinement across 12 datasets produce a CS gene signature with improved integrity

Corresponding gene expression and metabolite measurements are available only for *Bertilsson* (dataset ID 1). We thus implemented a bioinformatics strategy to identify a refined CS gene signature with improved integrity across all 12 datasets (2794 samples). We used the initial CS signature from *Bertilsson* as a starting point, and then used results from cancer samples across the other 11 datasets to nominate better gene candidates to replace genes from the initial CS signature (see Methods). When evaluating the new top 150 candidate genes, we found that 74 of the 150 genes in the initial CS signature had been replaced by new genes to improve integrity across all datasets (Figure 2C). Of note, the refined CS signature improved CMS for in both cancer and normal samples in all 11 additional datasets. This consistent improvement in normal samples (including normal samples from *Bertilsson*) affirms the relevance of the signature, since these samples were not used to generate or refine the CS signature. In addition, only a marginal reduction in CMS was observed for the cancer samples in *Bertilsson* (Figure 2D). We thus conclude that the refined CS signature (referred to as the CS signature from now) represents a more robust signature with improved integrity compared to the initial CS signature.

### The CS signature displays unique enrichment in prostate samples and negative correlation to stromal tissue compartments

Having established a CS signature of 150 genes associated with citrate and spermine secretion in the previous sections, we expanded our validation to additional datasets and sample types. For this analysis we used single samples Gene Set Enrichment Analysis (ssGSEA) [43], which is a method used to assess whether a specific gene signature is enriched or depleted in a single sample. Since citrate and spermine secretion is a highly specific function associated with prostate epithelium, higher ssGSEA scores in tissue samples from prostate compared to samples from other tissue types would confirm its functional relevance. Further, since prostate cancer samples are a heterogeneous tissue mixture of normal prostate epithelium, cancer and stroma, ssGSEA scores should be inversely proportional to the content of stroma tissue in the samples, since stromal cells are unable to secrete citrate or spermine.

We first compared ssGSEA scores for cancer and normal samples in the prostate specific *TCGA* dataset (dataset ID 5) to all cancer and normal samples profiled in the *TCGA-complete* resource (11093 samples from 33 cancer types, dataset ID 21). We observed strongly elevated ssGSEA scores for both prostate cancer and normal samples compared to cancer and normal tissues from other tissues (Figure 3A). We also tested the CS signature on average expression profiles for 53 normal tissues from the *GTEx* portal (dataset ID 22) (Figure 3B) and 1829 CAGE expression profiles from cell-lines and tissues in the *FANTOM* Consortium (dataset ID 23) (Figure 3C) where one tissue profile was from adult normal prostate. In both datasets, normal prostate tissue showed the highest ssGSEA scores.

**Figure 3:**
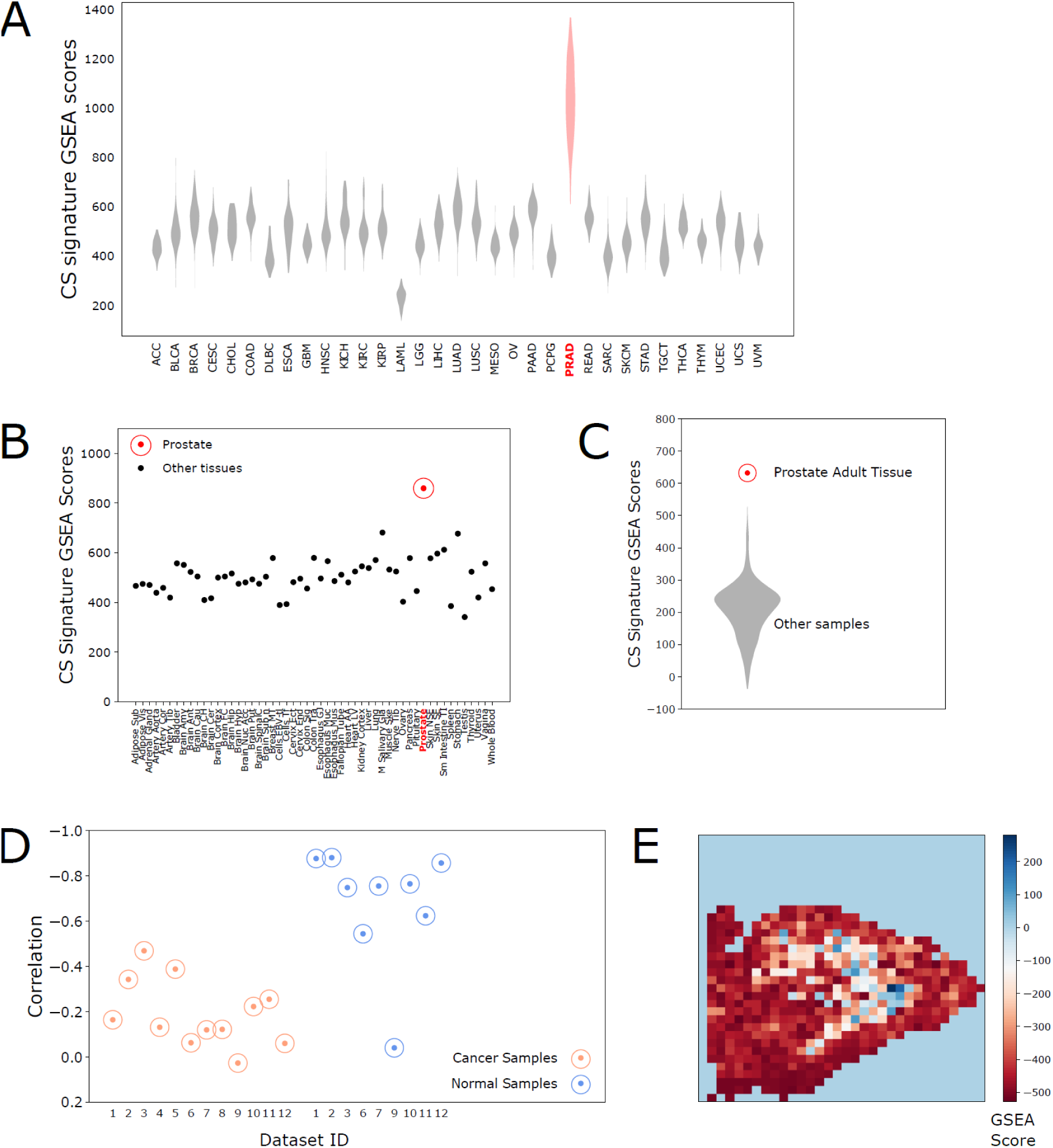
Validation of citrate – spermine (CS) gene signature. A) CS signature ssGSEA scores for 11093 cancer and normal samples in 33 tumor types from the TCGA-complete dataset. We have used standard TCGA cancer type abbreviations [https://gdc.cancer.gov/resources-tcga-users/tcga-code-tables/tcga-study-abbreviations] where PRAD is the abbreviation for prostate cancer. B) CS signature ssGSEA scores for 53 averaged human normal tissue profiles from the GTex dataset. C) CS ssGSEA signature scores for 1829 human cell-line and tissue samples, including one prostate adult tissue sample, from the FANTOM dataset. D) Correlations between CS signature and stroma signature ssGSEA scores for cancer and normal samples in prostate cancer tissue datasets 1-12. The negative correlations in the normal samples are stronger due to less confounding from cancer tissue. Data from Mortensen (dataset ID 9) contains laser dissected prostate normal epithelium and cancer, which probably lack significant amounts of stroma. E) CS signature ssGSEA scores on spatial transcriptomics data from tissue slice 3.2 in Berglund (dataset ID 20). The colorbar indicates enrichment of the ssGSEA score in each pixel. The corresponding pathological tissue image can be found for image 3.2 in Supplementary Information – Supplementary Figure 1b from Berglund et al. [33], and comparison shows a strong overlap of high and low CS signature scores with epithelial/lumen and stroma tissue compartments, respectively. Results from all 12 images in Berglund are shown in Supplementary Figure 2.

We next correlated the CS signature ssGSEA scores to a previously identified gene signature for stroma content in prostate tissue [44] in datasets 1-12. We observed a strong reverse correlation between CS and stroma signatures for normal samples in all datasets (Figure 3D) (with an exception for *Mortensen (*dataset ID 9) which contained laser dissected epithelium/cancer with minimal amounts of stroma). The correlation was weaker in cancer samples. This is expected, since prostate cancers vary in their ability to secrete citrate and spermine. The negative association between CS signature and stroma was also confirmed using spatial transcriptomics data from Berglund (dataset ID 20), where pixels with high and low ssGSEA scores overlaid regions with epithelium/lumen and stroma respectively (Figure 3E, Supplementary Figure 2).

In summary, the CS signature shows strong associations with expected properties of the prostate epithelium, which strengthens the biologically validity of the genes in the signature.

### The CS gene signature is gradually lost from normal samples through low grade, high grade and metastatic lesions

Having established the relevance of the CS signature to prostate citrate and spermine secretion, we next investigated how this signature associates with the various stages of prostate cancer aggressiveness. Low citrate concentrations have previously been associated with high-grade prostate cancer [4]. We identified nine datasets (1779 prostate cancer samples) where cancer samples had been classified as either high-grade or low-grade, or assigned a Gleason score - the standard form of prostate cancer grading. We found significant changes in CS signature ssGSEA scores between high- and low-grade cancers in all eight datasets where samples where Gleason score 4+3 was classified as high-grade (613 high and 737 low-grade samples), and six out of eight datasets where samples with Gleason score 4+3 was classified as low-grade (578 high- and 1258 low-grade samples) (Figure 4A, Table 1). The reduction in CS signature scores were not due to increased tumor fraction in high-grade samples (shown in Table 1. These results support the previous findings and suggest that the changes previously observed for citrate at metabolite level are accompanied by changes in gene expression. A further significant reduction in CS signature scores were observed in metastatic compared to cancer and normal samples in 8 datasets (157 metastatic samples) (Figure 4B, Table 2). Note that the lower CS-signature scores in normal samples compared to cancer samples is due to the high level of stromal tissue in normal prostate samples [8, 44] (Supplementary Figure 3). Overall, our results indicate that the genes associated with citrate and spermine secretion is upregulated in the normal epithelium, and then gradually downregulated with tumor progression from early low-grade lesions, through high-grade cancers and finally metastatic tumors. This trajectory is nicely illustrated by the results from *Tomlins* (dataset ID 18) (Figure 4C), where laser dissection has been used to purify stroma, normal epithelium, Prostate Intreapeithelial Neoplasia (PIN, an early pre-cancerous prostate lesion) cancer and metastatic tissues. In line with this trajectory, the PIN lesions show an average CS signature score level between the levels of normal epithelium and cancer. Though the ability of cells to secret citrate and spermine seems to be lost in metastatic samples, metastatic tissue from prostate cancer still contains traces of the CS gene profile compared to metastatic tissue originating from other organs (*Hsu*, dataset ID 19) (Figure 4D).

**Table 1:**
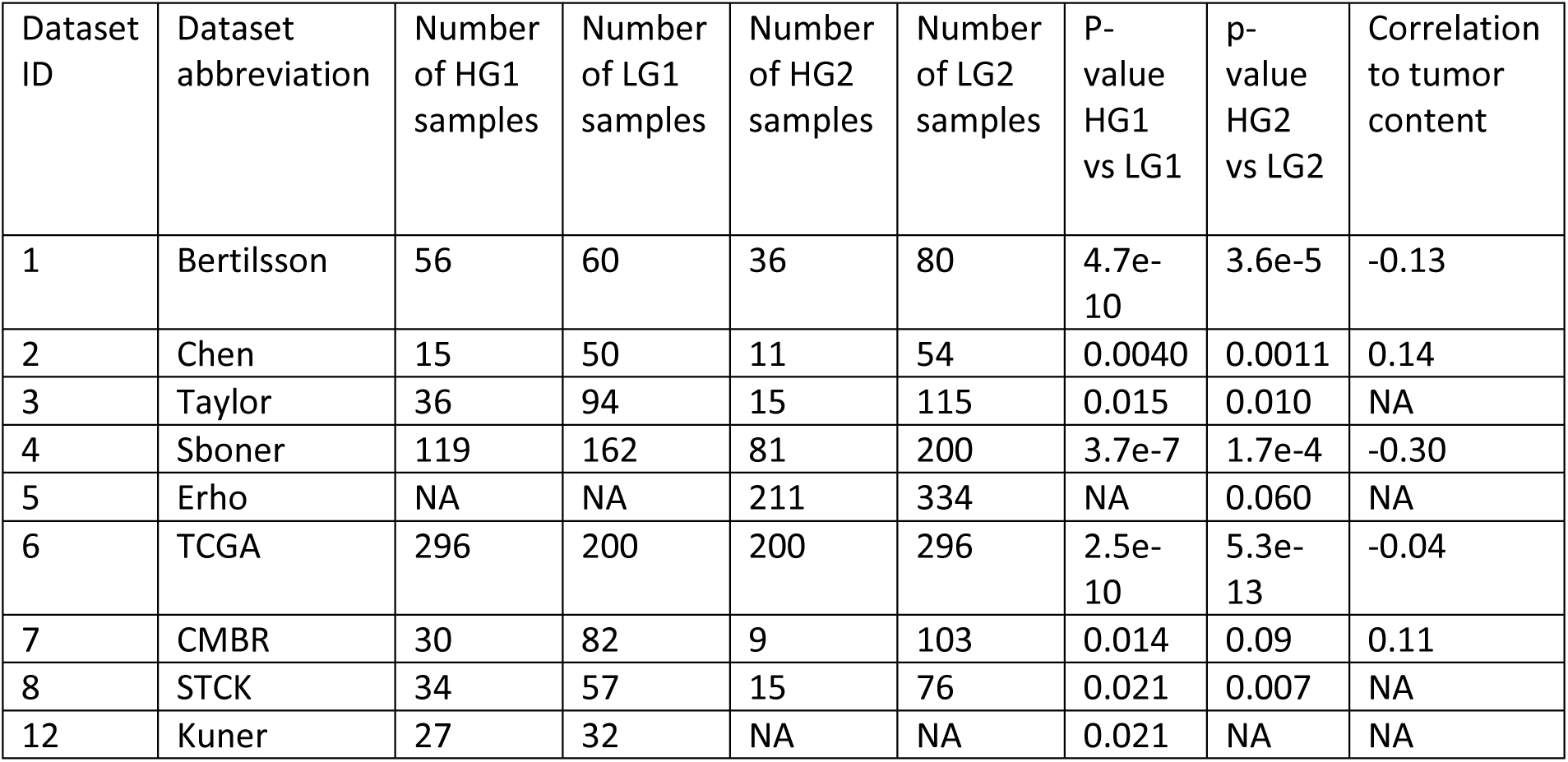
P-values from differential CS signature ssGSEA score analysis between high grade (HG) and low grade (LG) cancer samples in 9 datasets. Two comparisons are made: First HG1 vs LG1, where HG1 is classified as samples with Gleason score 4+3 or higher, and LG1 as samples with Gleason score 3+4 or lower. Second HG2 vs LG2, where HG2 is classified as samples with Gleason score 8 or higher, and LG2 as samples with Gleason score 7 or lower. The correlation of CS signature GSEA score to tumor content is also included when available, and show, in general, that decreased CS scores in high-grade samples is not due to increased tumor content in these samples.

**Table 2:** P-values from comparing CS signature ssGSEA scores from metastatic prostate cancer samples to cancer and normal samples and in 8 datasets.

| Dataset ID | Dataset abbreviation | Number of metastatic samples | Number of cancer samples | Number of normal samples | P-value metastatic vs cancer | P-value metastatic vs normal |
| --- | --- | --- | --- | --- | --- | --- |
| 3 | Taylor | 19 | 131 | 29 | 7.4e-15 | 2.8e-7 |
| 10 | Prensner | 12 | 78 | 38 | 3.4e-11 | 2.9e-10 |
| 13 | Aryee | 18 | NA | 21 | NA | 0.024 |
| 14 | Chandran | 25 | 65 | 81 | 8.8e-6 | 6.9e-6 |
| 15 | Cai | 29 | 22 | NA | 5.8e-13 | NA |
| 16 | Poisson | 13 | 12 | 16 | 1.5e-4 | 0.09 |
| 17 | Monzon | 21 | 10 | NA | 7.4e-11 | NA |
| 18 | Tomlins | 20 | 32 | 27 | 0.007 | 1.2e-10 |

**Figure 4:**
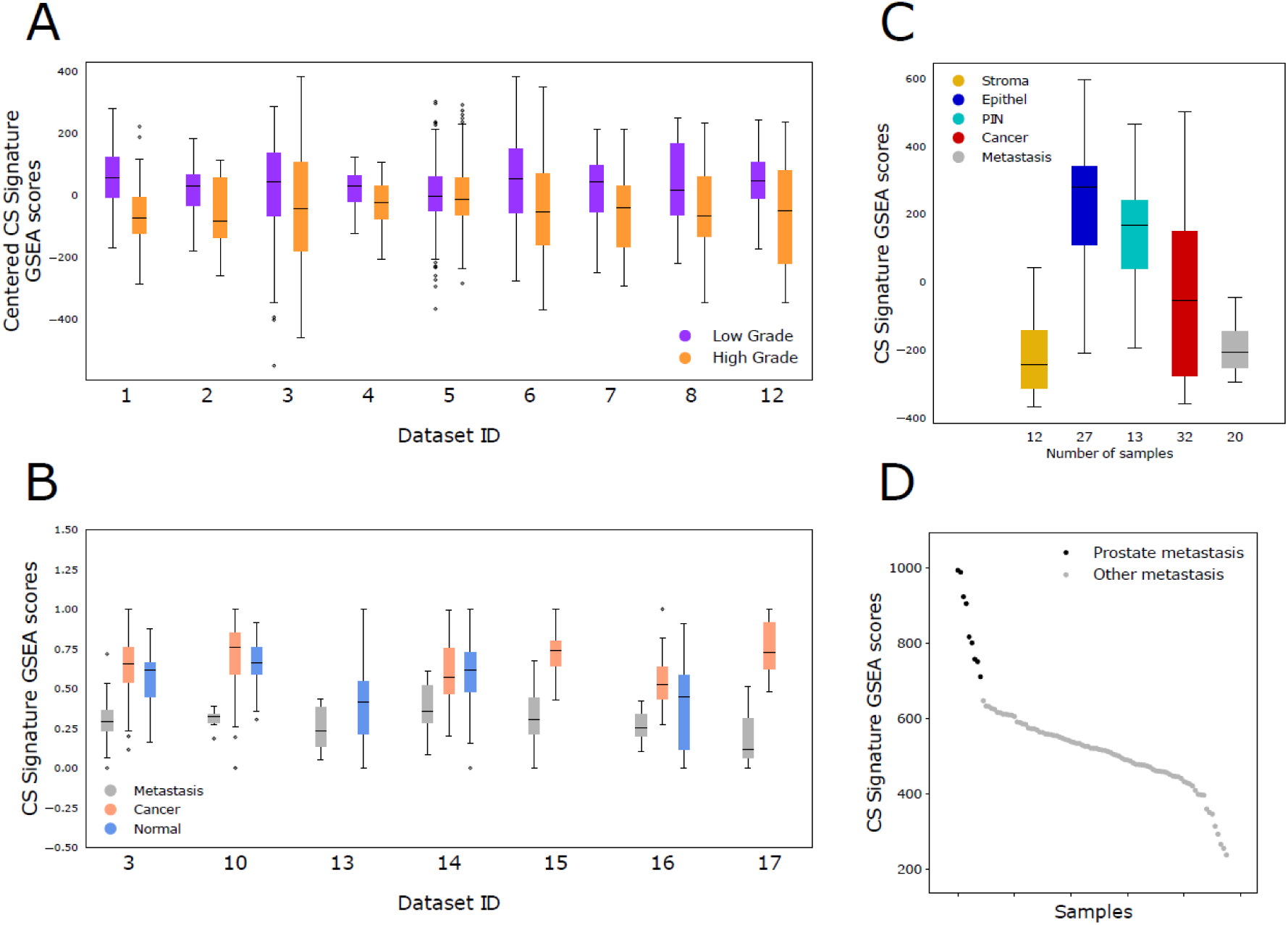
Citrate – spermine (CS) signature, tumor grade and metastasis. A) CS signature ssGSEA scores for high-grade (Gleason higher or equal to 4+3) and low-grade (Gleason less than or equal to 3+4) cancers in 9 datasets. The samples compared corresponds to the HG1 and LG1 groups in Table 1. The scores were centered in each cohort before plotting to visualize similarities between datasets better. B) CS signature ssGSEA scores for metastatic, cancer and normal samples in 7 datasets. The scores were centered and normalized to range 0-1 before plotting to visualize similarities between datasets better. The low scores in normal samples is due to the higher content of stroma in normal samples (Supplementary Figure 3). C) CS signature ssGSEA scores from five laser dissected prostate and prostate cancer tissue types from Tomlins (datasets Id 18). The GSEA scores illustrates the loss in CS secretion from normal epithelium through PIN, Cancer and finally Metastasis. The CS signature scores from metastatic samples are comparable to the non-secreting stroma tissue. D) CS signature ssGSEA scores in 96 metastatic samples (9 from prostate and 87 from other organs) from the Hsu (dataset ID 19). Metastatic samples from prostate origin are more enriched for the CS signature than metastatic samples originating from other sites.

### CS signature genes are enriched for genes with zinc-binding, mitochondrial and secretory functions

Gene Ontology analysis anchored the genes in the CS gene signature to the citrate secretory pathways from mitochondria towards the cell exterior via the ER and golgi. The analysis also highlighted several genes related to zinc-binding, in particular several metallothionines, and a role of branched chain fatty acid catabolism in upholding the normal prostate secretory function.

When examining gene functions on gene-by-gene basis using the NCBI-gene resource [ncbi.nlm.nih.gov/gene] (Supplementary File 3) we found that several genes in the CS signature were related to zinc-binding. Among these, six were Metallothioneins (*MT1G, MT1M, MT1F, MT1X, MT1E* and *MT2A*), one zinc-transporter (*SLC39A10*) and the two genes *AZGP1* (alpha-2-glycoprotein 1, zinc-binding) and *ANPEP* (alanyl aminopeptidase, membrane), the latter containing a consensus sequence known from zinc-binding metalloproteinases.

We then used DAVID [45, 46] and Enrichr [47, 48] to perform Gene Ontology (GO) analysis on the CS signature (Supplementary Figure 4). For both GO tools the most significant terms were related to *zinc/metal-ion binding* and *branched chain amino acid (Leucine, Isoleucine and Valine) degradation*. When inspecting the CS signature genes at NCBI, we noticed that many genes did not have any associated functional description, while at the same time having high prostate specific expression. A potential challenge for our GO analysis is that the prostate specific function of many of the genes in our CS signature is not known. We therefore applied GAPGOM, an alternative GO analysis tool where each gene is associated with functions based on consensus ontology terms for genes it is correlated with [49] (methods, Supplementary File 4). This analysis revealed strong gene ontology terms related to mitochondria, ER (endoplasmic reticulum), golgi, lysosome and exosome cellular components, which would fit well with a secretory path of citrate. When we performed Principal Component Analysis (PCA) to group genes according to their GO terms (see methods), the six metallothioneins formed a distinct cluster related strongly to zinc/metal-ion binding (Figure 5A, Supplementary File 4). In addition, the metallothioneins also showed strong associations with the mitochondrial respiratory complex and electron transport chain.

**Figure 5:**
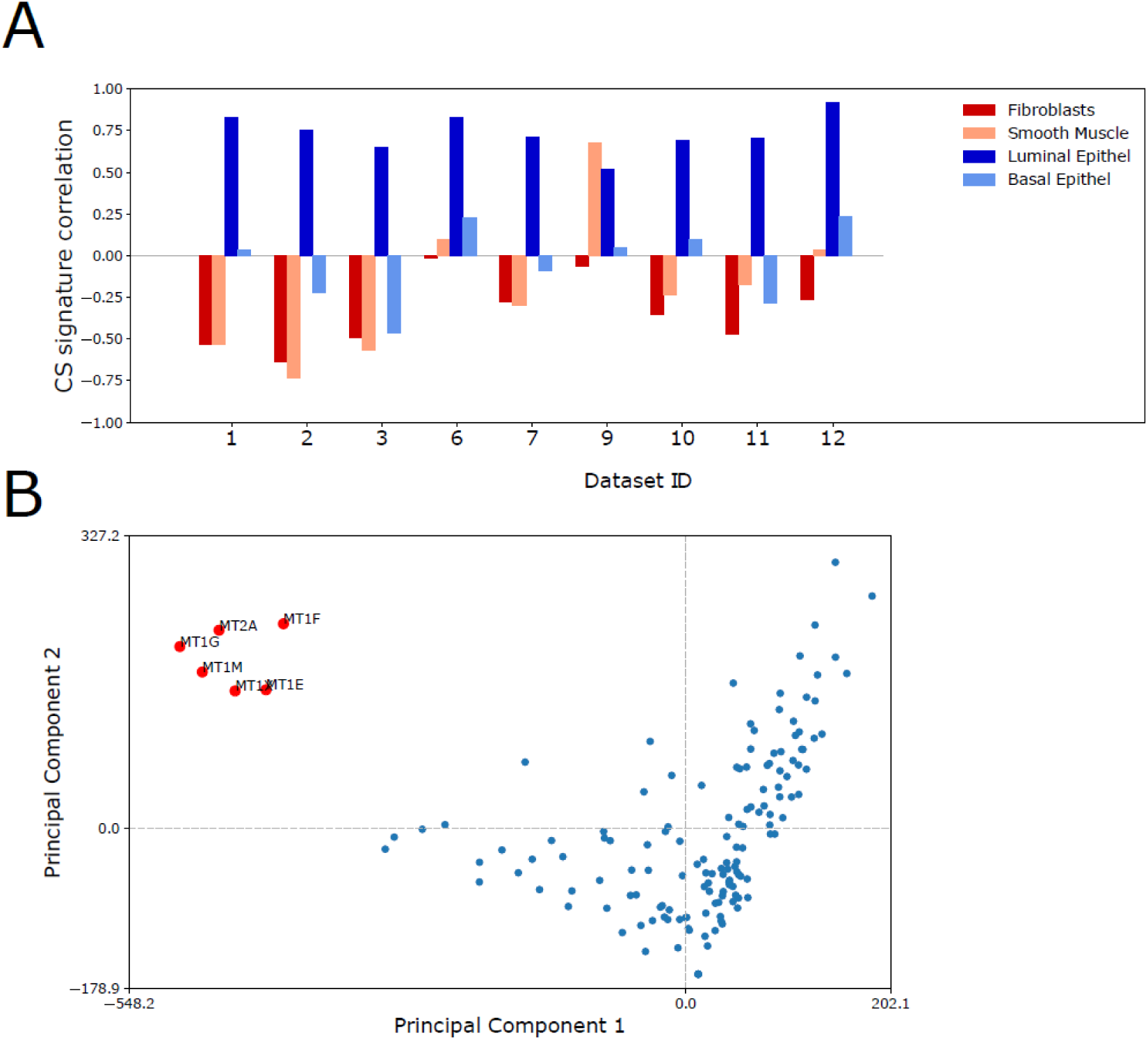
Genes and ontologies. A) Principal Component Analysis (PCA) for Gene Ontology terms associated with CS signature genes. The GO terms are based on a consensus analysis over datasets 1-12 (see methods). The only observable cluster is formed by the six Metallothioneins from the CS signature. B) Correlation of ssGSEA signature with four gene signatures from distinct cell types in a single cell sequencing dataset (basal and luminal from the prostate epithelium, fibroblasts and smooth muscle cells from stroma) [50] in 9 datasets with normal prostate samples. The CS gene signature correlates with luminal epithelial cells (the cell-type mainly responsible for secreting citrate and spermine), while the correlation to stroma tissue types is negative. Dataset 9 is from laser dissected epithelium, which lack stroma tissue.

We also performed network analysis on the 150 genes in the CS signature (Supplementary Figure 5), where the genes *(ALOX15B, RAB27A, ENDOD1, SLC45A3, NCAPD3, EHHADH, ACAD8, AFF3, NANS* and *YIPF1*) were identified as the top 10 hubs in the network. These hub-genes were all ranked among the top 20 in the CS-signature and had significant GO terms associated with branched chain fatty acid catabolism/degradation. Finally, we compared CS signature ssGSEA scores with cell-types specific gene signatures from a single-cell RNA-Seq study on normal prostate tissue [50]. The CS signature scores showed high correlation with prostate luminal cells (the cells mainly responsible for secretion), but not basal cells, and a negative correlation to stromal fibroblasts and smooth muscle cells (Figure 5B). Moreover, the luminal gene signature also included several of the metallothionein genes, corroborating the potential importance of these genes.

### The CS signature is depleted in prostate model systems

To reveal detailed function of the genes highlighted in the previous section will need further wetlab experiments. We therefore wanted to identify suitable model-systems where the possible role of these genes and their mechanisms could be further studied. Unfortunately, we found the identification of model systems to be a challenge.

We tested the CS signature on publicly available gene expression datasets from various model systems for prostate cancer using ssGSEA. In the *Prensner* (dataset ID 10), 58 expression profiles from cell-lines were analyzed together with tissue and metastatic samples, including the four most common prostate cancer cell-lines PC3, DU145, LNCaP, VCaP, and the normal prostate cell lines RWPE and PrEC. This unique collection of sample types makes the *Prensner* dataset an excellent reference for comparisons.

CS signature ssGSEA scores from the *Prensner* cell-lines were consistently at the low end, compared to both tissue and metastatic samples (Figure 6A), indicating that cell-lines derived from prostate do not secrete citrate and spermine. This observation was confirmed in *Taylor* (dataset ID 3), which also contained four cell-line samples (Supplementary Figure 6). When comparing different cell-lines, androgen responsive cell-lines (LNCaP and VCaP) generally have higher CS signature ssGSEA scores then androgen resistant (DU145 and PC3) cell-lines (Figure 6B). The overall highest CS scoring cell-line is MDA-Pca-2b (an androgen responsive cancer cell-line). Interestingly, normal prostate cell-lines (RWPE and PrEC) do not score higher than the cancer cell-lines. The relative differences in CS ssGSEA scores between the different cell-lines were highly stable, which was confirmed when assessing additional cell-line samples in *CCLE* (Cancer Cell Line Encyclopedia, dataset ID 24) (Figure 6C), *Søgaard* (Dataset ID 26) and *E-MTAB-2706* (dataset ID 25). (Supplementary Figure 7 and 8, respectively). Additionally, results from cohort *CCLE* and *E-MTAB-2706* show that androgen resistant prostate cancer cell-lines (DU145 and PC3) do not separate from cell-lines of other cancer origin in terms of CS signature GSEA scores (Figure 6C, Supplementary Figure 8).

**Figure 6:**
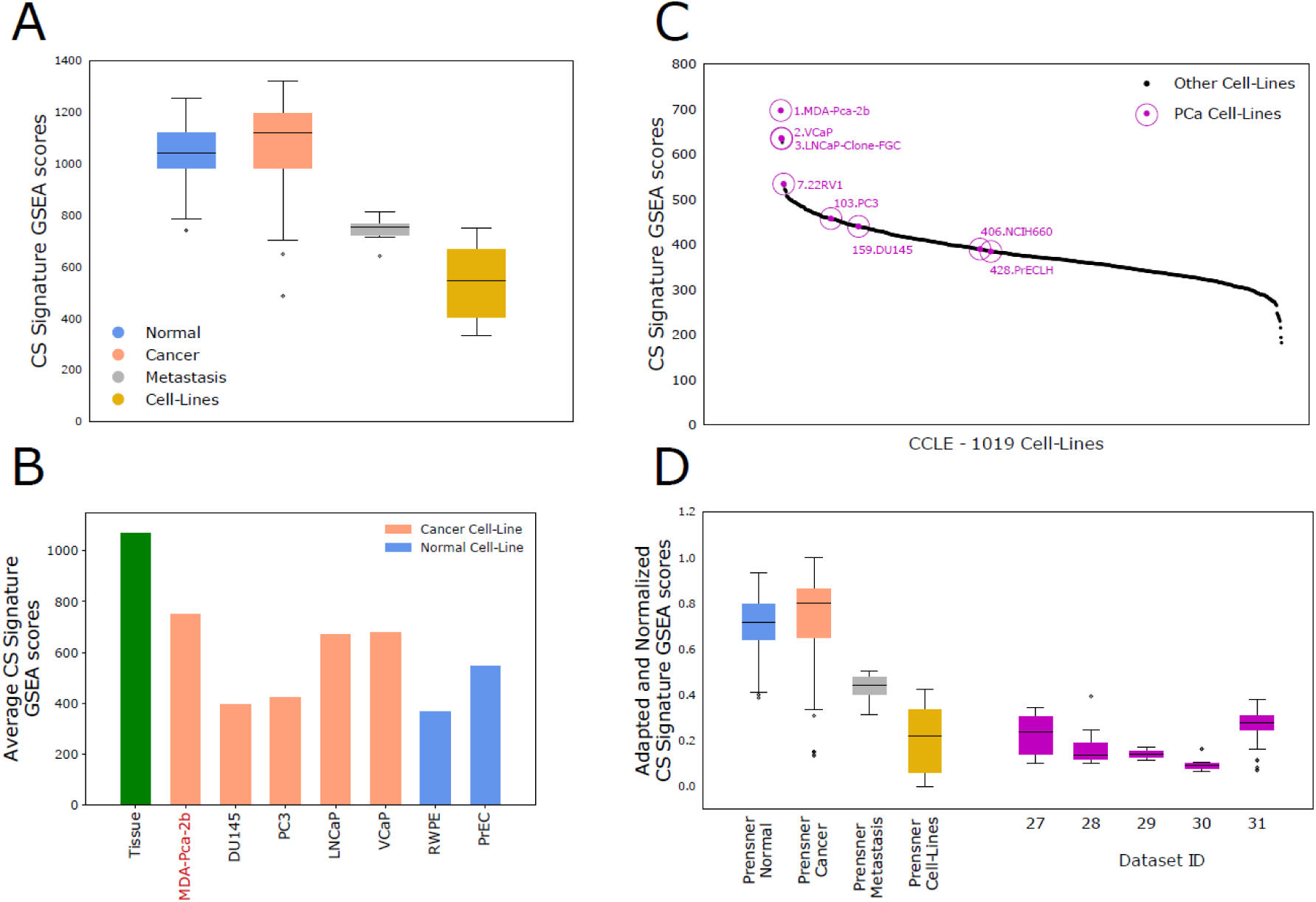
Model systems. A) CS signature ssGSEA scores for all sample types in Prensner (dataset ID 10), including 58 prostate normal and cancer cell-line samples. Cell-line samples, in general, have very low CS signatures scores. B) Average CS signature ssGSEA scores for different cell-types in Prensner. The ssGSEA scores were highly reproducable within samples from the same cell-type. Androgen responsive cell-types (MDA-Pca-2b, LNCaP and VCaP) have somewhat higher ssGSEA scores than androgen resistant cell-types (DU145 and PC3). Prostate normal cell-types are similar to cancer cell-types. Number of samples of each cell-type are: MDA-Pca-2b (1), DU145(10), PC3(2), LNCaP(7), VCaP(10), RWPE(9), PrEC(4). C) CS signature ssGSEA scores in 1019 cancer cell-types from CCLE (dataset ID 24). Androgen responsive prostate cancer cell-types score higher than cancer cell-types from other cancers, while androgen resistant cell-lines score similar to cell-types from other cancers. D) : CS Signature ssGSEA scores in four datasets with prostate and prostate cancer derived organoids and model systems (dataset IDs 27-30) and one dataset with prostate tumors from mouse models (Aytes, dataset ID 31). The ssGSEA scores were adapted and normalized to CS signature GSEA scores from Prensner. All model systems have consistently low GSEA scores, similar to prostate cancer cell-lines. In total 64 model system samples and 384 mouse tumor samples were analyzed. Dataset descriptions: **27**: 2D and 3D cultures derived from prostate cancer cell-line LNCaP (9 samples) and normal cell-line RWPE (9 samples), in total 18 samples. **28**: Organoids from primary prostate epithelial cells in mono and co-culture with stromal cells (7 samples), stromal cells in co-culture (4 samples) and macrodissected tumor tissue (2 samples), in total 13 samples.**29:** Cells from benign prostatic bulk (3+3 samples), basal (3+3 samples) and luminal (3+3 samples) tissue cultured in two different media, in total 18 samples. **30:** Mouse prostate organoids with WT (6 samples) and mutated (6 samples) gene SPOP, in total 12 samples. **31:** Prostate tumor tissue from mouse models with different genetic modifications, in total 384 samples. (See also Supplementary Figures 6-9).

To enable the comparison of ssGSEA scores between datasets, we implemented a method to adapt ssGSEA scores from different datasets to the *Prensner* dataset (see Methods). To demonstrate the utility of this implementation, we first adapted the CS ssGSEA scores from *Taylor* to *Prensner*, since both datasets included tissue samples (normal and cancer), metastatic samples as well as cell-lines. The CS ssGSEA scores where highly concordant down to the level of different cell-types between the two cohorts after adaptation (Supplementary Figure 9). We then used the adaptation strategy to compare datasets from organoids and mouse models (dataset IDs 27-31) to the *Prensner* dataset to investigate whether they would resemble tissue or cell-lines. None of the model-systems analyzed seemed to produce CS ssGSEA scores at levels similar to prostate epithelial tissue or low-grade primary cancer tissue. Instead their CS scoring range typically compare to the cell-lines from *Prensner* (Figure 6D). Overall, these results call attention to the research challenge for current prostate model systems to recapitulate the function of normal prostate epithelium in terms of citrate and spermine secretion.

## Discussion

### Expanding the view of zinc-binding proteins and zinc-transporters in the prostate

Six metallothioneins were identified as part of the CS gene signature in this study and constituted a functionally distinct cluster particularly associated with zinc-binding. Metallothioneins are low molecular weight, metal binding proteins localized to the Golgi, and have different expression in normal prostate and prostate cancer [51, 52]. In addition, expression levels can change in response to zinc-stimuli in prostate cell-lines [53]. It has also been speculated that metallothioneins can affect mitochondrial function, since they are small enough to enter the mitochondrial membrane bilayer carrying zinc [54]. This suggestion fits with the GO associations of metallothioneins to mitochondria, respiratory complex and the electron transport chain. Otherwise, the other potential zinc-binding genes discovered in this study have not been studied previously in the context of citrate accumulation and secretion in the prostate.

The zinc-transporter SLC39A1 (ZIP1) was not a part of the CS-signature, though this gene has previously been shown to be important for zinc-homeostasis in prostate cells. There could be several reasons for this discrepancy. First, this study only measures genes at the transcript level, while it is possible that SLC39A1 is regulated at the protein level [6]. However, expression levels from genes displayed in NCBI-gene resource show that SLC39A1 is not prostate specific, but expressed in most tissues. Though SLC39A1 has the ability to transport zinc within prostate tissue, it may not be the main determinant of prostate specific zinc regulation *in vivo*. In this context is should also be noted that none of the metallothioneins displayed prostate specific expression in the NCBI-gene database. Of the zinc transporters, most prostate specific expression is observed for SLC39A2, SLC39A6, SLC39A7 and SLC39A10, where SLC39A10 was part of the CS signature. SLC39A6 and SLC39A7 (though not part of the signature) were also positively correlated with citrate in our analysis, while SCL39A1 and SLC39A2 showed no correlation. Of the potential zinc-binders, AZGP1 show the most prostate-biased expression, but the function of this gene is unknown. Overall, more targeted research experiments are needed to identify the genes that control zinc-levels in the prostate, and how they perform their function.

Previous studies have shown that prostate epithelial cells can utilize glucose and aspartate as the carbon sources for citrate production. The gene *GOT2* (*glutamic-oxaloacetic transaminase 2*, also named *mAAT*-*mitochondrial aspartate aminotransferase* in previous reports) was shown to be responsible for the conversion of aspartate to oxaloacetate and citrate in the mitochondria. *GOT2* was a part of the CS signature, in addition to *SMS* (*spermine synthase*), and both *GOT2* and *SMS* showed high tissue-expression in prostate according to NCBI-Gene. For these two genes our results fit with previous mechanistic knowledge [9, 55].

Very little is known about the role of branched chain amino acid catabolism for normal prostate function. All of the three amino acids leucine, isoleucine and valine are upregulated in prostate cancer [12], indicating their relevance for cancer transformation, and this fits with our finding that the degradation of these amino acids is important to maintain normal prostate function. Branched chain amino acids can possibly work as precursors for citrate production [56], and leucine can act as a sensor for mTOR-pathway activation [57], which is generally regarded as an important pathway in prostate cancer development [58].

### Expression of CS signature genes are with prostate cancer progression

There has been a discussion on whether the reduced citrate levels observed in prostate cancer are merely a result of a reduction in luminal glands leading to reduced amounts of prostatic fluid [59]. However, our results clearly show that changes in citrate and spermine levels are accompanied by changes at the gene expression level, and this study supports the hypothesis that changes in genes responsible for citrate and spermine secretion are important for the cell transformations leading to cancer. Further research into these mechanisms can lead to new discoveries and treatment targets in the management of prostate cancer.

### The challenge of finding a model system for normal prostate

We could not find CS signature enrichment comparable to tissue samples in any of the cell-lines and model systems we investigated. Androgen responsive cell-lines seem to maintain some traces of prostate specific functions compared to other model systems, but their CS signature enrichments are far below that of *in vivo* tissue and indicate that most parts of the prostate-lineage specificity is lost. Thus, these observations question the relevance of these model systems for studying normal prostate function and transitions from normal cells to prostate cancer cells. The secretory function can possibly be triggered by proper stimuli like dihydrotestosterone (DHT), testosterone or prolactin [60-62]. For example, LNCaP (but not PC3) cells were able to secrete citrate when stimulated by DHT [61]. However, it was also observed that the rate of citrate consumption by the TCA cycle increased proportionally in the same experiment [61]. Since these experiments were performed before the era of transcriptomics, it would be interesting to repeat these experiments with accompanying transcriptomics analysis to identify the genes that change during such stimuli.

### Signature refinement produces more robust gene signatures with improved integrity

In this manuscript, we have argued that improved gene signature integrity (assessed by CMS) can be achieved by integrating data from multiple prostate cancer datasets. The aim was to remove noise and produce better and more robust gene signatures in terms of biological interpretation. We found several evidences that more robust signatures were achieved by integration of datasets. First, we observed that the refined signature improved CMS scores compared to the initial signature in independent normal prostate samples (Figure 2D). Second, when we used the initial gene signature for ssGSEA in the *Hsu* dataset, we observed no clear separation between prostate and other metastatic samples (Figure 4D, Supplementary Figure 10). Third, in the *FANTOM* dataset, the one Prostate Adult Tissue sample were only ranked as the highest scoring sample after signature refinement, but not when the initial signature was used (Figure3D, Supplementary Figure 11).

The refinement procedure generated an improved CS signature ssGSEA scores which was particularly pronounced for prostate samples. We also tested the effect of refinement ion gene expression data from other tissue types in the *TCGA-complete* dataset (33 cancer and 22 normal tissue types). Other cancer types showed both elevated and decreased GSEA scores after refinement, and elevations were consistently lower than in prostate samples (Supplementary Figure 12).

The best CS signature ssGSEA score improvement after refinement were also observed particularly for the metabolites citrate and spermine. For this test, we redid the procedure to create initial and refined signatures for all metabolites and lipids in the metabolite data from *Bertilsson* (23 metabolites and 17 lipid signals). We then calculated ssGSEA scores in all cancer and normal tissue types in the *TCGA-complete* dataset for each metabolite/lipid signature (Supplementary Figures 13-16). The most elevated ssGSEA scores were for citrate and spermine in prostate compared to other tissue types, while other metabolites showed both elevated and decreased ssGSEA score for other tissue types compared to prostate. This observation was similar for both cancer and normal tissue samples. Thus, the effect of refinement was highest for citrate and spermine specifically in the prostate. Interestingly, we noted that other metabolites/lipid signatures were elevated in prostate, particularly putrescine (a precursor for spermine in the polyamine pathway) a few lipid signals and glucose, indicating prostate specific regulation of other metabolites than just citrate and spermine. These observations were confirmed in the *FANTOM* dataset (dataset ID 23) (Supplementary Figure 17).

In summary, we conclude that the identification followed by refinement procedure used in this study created a robust and unbiased gene signature biologically relevant for in vivo spermine and citrate secretion in the prostate. In a wider perspective, the module approach may represent a general strategy to find robust genesets related to other –omics data (for example metabolomics, proteomics or lipidomics), as long as these -omics data are accompanied with gene expression data.

## Conclusion

The genes and mechanisms governing normal prostate function, in particular the ability to accumulate and secrete large amounts of citrate and spermine, are mostly unknown. We have used bioinformatics on an extensive collection on prostate datasets to discover genes relevant for secretion of citrate and spermine in the prostate. Based on this we have created a 150 gene signature which can be used to assess the degree of citrate and spermine secretion in a single sample. The bioinformatics approach enabled us to validate and explore the gene signature in a large number of contexts based on public data, including normal vs cancer enrichment, spatial colocalization on tissue, tumor grade, metastasis and different model systems. The employed approach is generalizable to other types of data when measured together with gene expression. The signature showed an inverse association to cancer progression from normal through low and high grade to metastasis. The genes in the signature were enriched for zinc-binding, mitochondrial function, respiratory complex, secretion through the ER – golgi - lysosome – exosome pathway and branched chain amino acid catabolism, pointing to known, suggested and novel mechanisms for normal prostate function. The lack of suitable model systems is a challenge, and need to be established to study these findings further.

## Methods

### Datasets

A list of all datasets with description and references are given in Supplementary File 1

### Module refinement procedure

A schematic representation of the procedure with stages 1-3 is shown in Figure 7A. The procedure make use of the following abbreviations: **M**=Module, **G**=several genes, **g**=single gene, **D**=Dataset, **P**=gene expression Profile, **TB**=Table, **av**=average, **in**=initial, **rf**=refined, **c**=candidate, **m**=missing, **cr**=correlation, **ic**=intra-correlation, **CMS** = Correlation Module Score (CMS), **MGCR** = Module Gene Contribution Score (MGCR).

**Figure 7:**
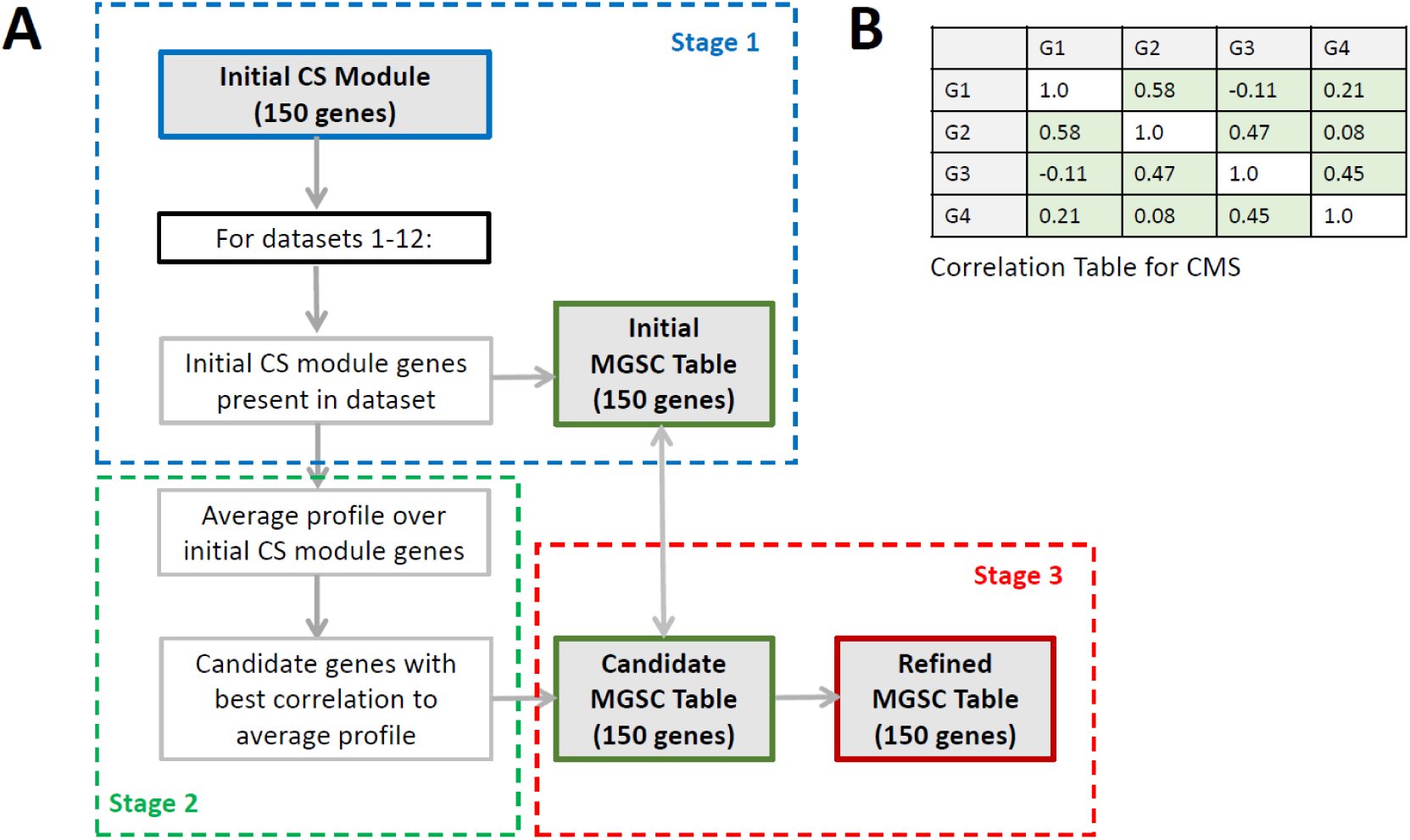
Module refinement procedure and Correlation Module Score (CMS) A) Flowchart of the module refinement procedure. Each stage 1-3 are explained in the main text. B) Example of a correlation table used to calculate the Correlation Module Score (CMS). This gene module has four genes (G1-G4) and a CMS of 0.32.

### Correlation Module Score (CMS)

To assess how well the correlations between genes in a module (or geneset) is preserved for all (or a subset of) samples in a particular dataset, we created the Correlation Module Score (abbreviated CMS). The CMS for a module in a dataset is calculated by first creating a table of intra-correlations between expression profiles for all genes in the module for that dataset (Figure 7B). Here the expression profile for a gene is the vector consisting of gene expression values over all samples in that dataset. For a 150-gene module, this will create a table with 150*150 = 22500 correlation values (a table for a hypothetical 4-gene module is shown in Figure 7B). The CMS is then calculated as the average over all correlations included the table (but excluding self-correlations). A high CMS represents a high integrity of the gene module in the dataset, indicating the preservation of strong correlations between the module-genes in the datasets. Likewise, a low CMS would indicate weak relations between the genes in the module, and the loss of gene module integrity. We further assume if a gene module has a high CMS in a dataset, it would indicate that the biological process the module represents is important or active for samples in this dataset. In this study the datasets are patient cohorts with normal and cancer tissue samples from the prostate.

### Refinement Procedure - Stage 1

The purpose of Stage 1 in the module refinement procedure is to find the contribution of each gene in the initial CS module from *Bertilsson* to the overall CMS for the module. To represent this contribution we introduce the Module Gene Contribution Score (MGCS), which is the average over all correlations between that gene and each of the other genes in the module (or, the overall contribution of that gene to the overall CMS for the module). In Figure 7B this will be the average of the rows (excluding the diagonal), where each average value represent the MGCS score for a gene. A high MGCS score will indicate that the expression profile of the gene fits better with expression profile of other genes in the module, while a low MGCS score indicates a gene expression profile that deviates from other genes in the module.

#### Procedure

S1: For each dataset **D** (dataset ID 1-12 with normal and cancer tissue samples from the prostate): Find the genes from the initial CS module, **in-M-G**, present in the dataset **D, in-M-G-D**

Calculate the intra-correlation table, **ic-TB-D**, for the initial CS module gene profiles, **in-M-G-P-D**, in dataset **D**

Calculate the corresponding CMS score, **CMS-D**, from the intra-correlation table, **ic-TB-D**, in dataset **D**

Calculate the MGCS score for each initial CS gene **in-g** in **CMS-D, in-g-MGSC-D**

For missing initial CS module genes, **m**-**in-M-G-D**, in dataset **D**: Set the **in-g-MGSC-D** to a *dataset specific missing gene constant* equal to the **CMS-D** multiplied by 0.5

Store the **in-g-MGCS-D** for each initial CS module gene in each dataset **D** in an *Initial MGSC table*, **in-MGCS-TB** for comparison in Stage 3.

Store MGSC for missing genes in each cohort, **m-MGSC**, for **Stage 2**.

### Refinement Procedure - Stage 2

The purpose of Stage 2 is to calculate, for all candidate genes **c-g** present in any one of the 12 datasets **D**, a score which describes how well each candidate gene expression profile fit with the expression profiles of genes in the initial CS module. This is done by comparing each candidate gene expression profile **c-g-P-D** to the average expression profile, **in-M-G-P-D-av**, of the genes in the initial CS module for each dataset **D**. We regard any gene present in any of the 12 datasets **D** as candidate genes.

#### Procedure

Define candidate genes, **c-G**, by the union of all genes present in any dataset **D**

For each dataset **D**:

Find the genes from the initial CS module, **in-M-G**, present in dataset **D**

Calculate the average expression profile, **in-M-G-P-D-av**, over expression profiles for all initial module genes **in-M-G-P-D** in dataset **D**

For each gene **g** present in dataset **D**:

Calculate the correlation between the expression profile of gene **g** and the average profile **in-M-G-P-D-av** for dataset **D**

For genes not present in this dataset, the correlation is set to the *cohort specific missing gene constant* in **m-MGSC** from **Stage 1**

Store all correlations in a table, **cr-TB**, for evaluation in **Stage 3**

### Refinement Procedure - Stage 3

The purpose of Stage 3 is to identify the genes that correlates best with genes in the initial CS module over all 12 datasets, but that were not a part of the initial CS module to begin with. From this analysis we select the 150 best candidates, and compare their MGCS to the MGSC for the genes in the initial CS module from **Stage 1**. If the MGSC for a new candidate gene is higher than the MGSC for an initial CS module gene, the initial gene is replaced by the candidate gene. The final output is a refined CS module, where the robustness of gene correlations across all 12 prostate cancer datasets are taken into consideration in the selection of genes for the module. This refined module is defined as the CS gene signature.

#### Procedure

Exclude initial CS module genes **in-M-G** from the correlation table **cr-TB** from **Stage 2**

Calculate the average correlation, **c-g-cr-av**, for each of the remaining candidate genes over all 12 datasets in **cr-TB**.

Sort genes in correlation table **cr-TB** by average correlation **c-gr-cr-av**.

Select 150 candidate genes, **c-g-150**, with the highest average correlation, **c-gr-cr-av**

For each dataset **D**:

For each of the top 150 candidate genes, **c-g-150**:

Calculate the average correlation of gene **c-g-150** to all genes from the initial CS module, **in-M-G-D**, and use the average correlation as the candidate gene MGSC, **c-g-MGSC-D**. This is a measure of the fit for the candidate gene, **c-g-150**, to the initial CS module in dataset **D**.

For candidate genes not present in dataset **D**, the MGSC is set to the *cohort specific missing gene constant* in **m-MGSC** from **Stage 1**

Combine the candidate MGSC table **c-g-MGSC-D** with the initial MGSC table **in-g-MGSC-D** from **Stage 1**. The combined table, **comb-TB**, will contain 300 genes, 150 from the initial CS module and 150 new candidates.

Calculate the average MGSC score, **g-MGSC-av**, for each gene over all 12 datasets for the combined table **comb-TB**.

Sort genes by average MGSC score, **g-MGSC-av**.

Select the 150 genes with the highest average MGSC score, **g-MGSC-av**, as genes for the refined CS module, **rf-G-M**.

The number of genes from the initial CS module present in each of the 12 datasets are shown in Supplementary Table 1.

### Single sample Gene Set Enrichment Analysis (ssGSEA)

Single sample Gene Set Enrichment Analysis (ssGSEA) was performed by sorting the genes in each sample according to their expression level in descending order. Then a ssGSEA score was calculated for each sample implementing a previously published algorithm [43]. We did not perform centering and normalization of the expression values for ssGSEA in this study. This is because citrate and spermine secretion, as well as the content of stroma tissue in the sample, are dominating biological features of these samples where differences should be visible at absolute expression level. However, we observed good correlations (0.86 – 0.97) for ssGSEA scores based on normalized and absolute expression values. A few of the datasets were only presented with centered expression values in the public resource, and for these we used centered values for ssGSEA. For the spatial transcriptomics data from *Berglund* (dataset ID 20), we did use centered and normalized expression for ssGSEA in each pixel. This was necessary due to many pixels with a high number of unexpressed genes, and/or very few genes with abundant expression.

### Adapted and normalized ssGSEA

The magnitude of GSEA scores depend on both the size of the gene set used, and the number of total genes in the ranked list input to the GSEA calculations. To be able to compare ssGSEA scores from *Prensner* to other datasets, we adapted the ssGSEA by only using the genes shared between the datasets in each comparison. In this way, both the number and identity of genes, as well as the size of the geneset will be identical for both datasets. For the comparison of several datasets in Figure 4D, we also normalized the GSEA scores in *Prensner* to 0-1 scale for each comparison. The adapted and normalized score calculated in each comparison were highly correlated (average 0.99), and had a small standard deviation of the mean (0.004), and we conclude that they are comparable between the datasets.

### GO analysis

The GO analysis was based on gene annotation generated with the GAPGOM tool [49] [PMID: 30567492, GAPGOM in Bioconductor]. GAPGOM will predict gene annotation for a target gene by identification of well-annotated genes showing correlated expression pattern with the target gene across several experiments, and then estimate a predicted annotation for the target gene as a consensus over the co-expressed genes, based on the hypothesis that co-expressed genes may be involved in similar or related processes. This prediction of annotation terms may facilitate a richer functional annotation of genes, in particular for genes where there is a lack of experimental annotation data.

Since GAPGOM only allows us to do annotation prediction one gene at a time, GAPGOM was used in conjunction with Snakemake [https://snakemake.readthedocs.io/en/stable/project_info/citations.html] to create a software pipeline. This pipeline was used to predict GO term annotation for all 150 genes in the CS-signature on each of dataset 1-12. Since GO annotation consists of three different ontologies (MF, BP, CC), each prediction is done separately for each. This then outputs a list for each gene in each dataset for each GO ontology, in total 1800 lists. Each list contains one or multiple GO-term predictions, with the following information for each prediction; GO-ID, Ontology, P-value, FDR/q-value (Bonferoni normalized P-value), The description of the GO term, and the used correlation method for the prediction. For each list, only GO terms with a q-value > 0.05 were selected. This generated a total of 21 080 GO terms for all lists, where 5354 GO terms were unique. A table was created with the number of genes as rows and the number of unique GO-terms as columns. For each gene and GO-term, we summed –log10 q-values from all datasets, producing an overall score for each GO-term and gene. We made three tables, one for all 150 genes, one for the six Metallothioneins, and one for the 10 network Hub-genes.

### Network analysis

Correlation-based networks of the CS signature genes for each dataset are created as follows: The 150 genes are represented as nodes in the network and the pairwise Pearson correlation between genes represents the interactions between nodes. To reduce the complexity and highlight the most powerful interactions, only the 20 strongest outgoing links (in absolute Pearson correlation) from each node are kept. In this way, the most central nodes will have 20 outgoing links and up to 130 ingoing links from other nodes, while some of the non-central nodes will only have 20 links. By calculating the node-degree (number of links to other nodes) for each node, genes with high node-degree reflects central genes for driving the biological processes [63]. This network construction is performed on cancer samples for all 12 datasets, and the 10 genes with the highest mean degree across the datasets are considered as the top 10 hub genes.

## Supplementary Material

**Supplementary File 1: Overview of datasets with references used in this study**.

**Supplementary File 2: Supplementary tables and figures**

**Supplementary File 3: CS signature genes and manually curated functional annotation**

**Supplementary File 4: Top 20 GO terms for all CS signature genes, six Metallothioneins and 10 network Hub-genes in the categories Biological Process (BP), Molecular Function (MF) and Cellular Component (CC)**.

## Acknowledgements

This works supported by the Liaison Committee between the Central Norway Regional Health Authority (RHA) and the Norwegian University of Science and Technology (NTNU) to [MBR]; the Norwegian Cancer Society [100792-2013] to [TFB], PhD position from Strategic funding ISB, Norwegian University of Science and Technology (NTNU) to [MKA], PhD position from Enabling Technologies, Norwegian University of Science and Technology (NTNU) to [KR]. The technique for fresh frozen tissue biobanking and cylinder extraction for reference E-MTAB-1041 was developed by Biobank1, St.Olavs Hospital, Trondheim, Norway. Funding support for MPC Transcriptome sequencing to identify non-coding RNAs in prostate cancer was provided through the NIH Prostate SPORE P50CA69568, R01 R01CA132874, the Early Detection Research Network (U01 CA111275), the Department of Defense grant W81XWH-11-1-0331 and the National Center for Functional Genomics (W81XWH-11-1-0520). The results shown here are in part based upon data generated by the TCGA Research Network: http://cancergenome.nih.gov/. The Genotype-Tissue Expression (GTEx) Project was supported by the Common Fund of the Office of the Director of the National Institutes of Health, and by NCI, NHGRI, NHLBI, NIDA, NIMH, and NINDS. The data used for the analyses described in this manuscript were obtained from the GTEx Portal on 11/23/2017.

## Author Contributions

MBR, MBT and HB conceived the idea and developed the concept. MBR, MBT and HB performed data curation. MBR developed methods and performed overall analysis. FD and CM developed and performed the GAPGOM analysis. MH performed network analysis. MBR, MBT and TFB acquired funding. MBR, MBT, SB, FD and TFB performed supervision and provided intellectual input. MBR, MBT, SB, HB, FD and TFB prepared the original draft. MBR prepared figures. All authors wrote and reviewed the manuscript.

## Ethical statement

The use of human tissue material and clinical data from the *Bertilsson* cohort was approved by the Regional Committee for Medical and Health Research Ethics (REC) for Central Norway, approval no 4-2007-1890. All experiments were performed in accordance with relevant guidelines and regulations. Informed consent was obtained from all participants. RNA-Seq data from the *Prensner* cohort was approved through dbGap (project #5870), and data were downloaded and stored according to the provided security requirements. All other data were downloaded from freely available and publicly open resources

