## Supplementary_File_1 for "The genes controlling normal function of citrate and spermine secretion is lost in aggressive prostate cancer and prostate model systems"

| Dataset ID | Dataset Abbreviation | Description | Dataset source | Reference |
|---|---|---|---|---|
| 1 | Bertilsson | 156 prostate tissue samples<br>(116 cancer and 40 normal)<br><br>129 of these samples (95 cancer and 34 normal) with concordant metabolite and gene expression measurement. | Array Express:<br>E-MTAB-1041 | [1] |
| 2 | Wang | 136 prostate tissue samples<br>65 cancer and 71 normal | GEO: GSE8218 | [2, 3] |
| 3 | Taylor | 160 prostate tissue samples<br>(131 cancer and 29 normal)<br>19 prostate metastatic samples<br>4 cell-line samples (DU145, PC3, LNCaP, VCaP) | GEO: GSE21034 | [4] |
| 4 | Sboner | 281 prostate cancer samples | GEO: GSE16560 | [5] |
| 5 | Erho | 545 prostate cancer samples | GEO: GSE46691 | [6] |
| 6 | TCGA-PRAD | 549 prostate tissue samples | https://portal.gdc.cancer.gov/repository | https://www.cancer.gov/tcga |

| | | | | |
|---|---|---|---|---|
| | | (497 cancer and 52 normal) | | |
| 7 | CMBR (Cambridge) | 186 prostate tissue samples<br>(112 cancer and 74 normal) | GEO: GSE70768 | [7] |
| 8 | STCK (Stockholm) | 94 prostate cancer samples | GEO: GSE70769 | [7] |
| 9 | Mortensen | 50 prostate tissue samples<br>(36 cancer and 14 normal)<br>Laser dissected tissue | GEO: GSE46602 | [8] |
| 10 | Prensner | 116 prostate tissue samples<br>(78 cancer and 38 normal)<br>12 prostate metastatic samples<br>58 cell-line samples | dbGap: phs000443.v1.p1 (tissue and metastatic samples)<br>GEO: GSE31728 (Cell-lines) | [9] |
| 11 | Penney | 424 prostate tissue samples<br>(264 cancer and 160 normal) | GEO: GSE62872 | [10] |
| 12 | Kuner | 98 prostate tissue samples<br>(59 cancer and 39 normal) | GEO: GSE32571 | [11] |
| 13 | Aryee | 21 normal prostate samples<br>18 prostate metastatic samples | GEO: GSE38241 | [12] |

| | | | | |
|---|---|---|---|---|
| 14 | Chandran | 145 prostate tissue samples<br>(65 cancer and 81 normal)<br>25 prostate metastatic samples | GEO: GSE6919<br>Data from GPL8300 Platform | [13, 14] |
| 15 | Cai | 29 prostate cancer samples<br>22 prostate metastatic samples | GEO: GSE32269 | [15] |
| 16 | Poisson | 28 prostate tissue samples<br>(12 cancer and 16 normal)<br>13 prostate metastatic samples | GEO: GSE8511 | [16] |
| 17 | Monzon | 10 prostate cancer samples<br>21 prostate metastatic samples | GEO: GSE6752 | [13] |
| 18 | Tomlins | 84 prostate tissue samples<br>(32 cancer, 13 PIN lesions, 27 normal epithelium and 12 normal stroma)<br>20 prostate metastatic samples<br>Laser dissected tissue | GEO: GSE6099 | [17] |
| 19 | Hsu | 96 metastatic samples from various cancers<br>(9 from prostate metastasis) | GEO: GSE18549 | [18] |

| | | | | |
|---|---|---|---|---|
| 20 | Berglund | Spatial transcriptomics on 12 slices from prostate cancer and normal tissue | http://www. spatialtranscriptomicsresearch.org/ | [19] |
| 21 | TCGA-complete | 11093 samples from 33 cancer types | https://portal.gdc.cancer.gov/repository | https://www.cancer.gov/tcga |
| 22 | GTex | Median profiles for 53 normal tissue types | https://gtexportal.org/home/datasets | https://gtexportal.org/home/ |
| 23 | FANTOM | 1829 samples from various human cell-lines and tissues | fantom.gsc.riken.jp/5/data/ | [20, 21] |
| 24 | CCLE | 1019 cell-line samples from various cancer types | https://portals.broadinstitute.org/ccle/data RNA-Seq data | [22] |
| 25 | E-MTAB-2706 | 622 cell-line samples from various cancer types | Array Express: E-MTAB-2706 | [23] |
| 26 | Søgaard | 36 samples from 3 prostate cancer cell-lines: DU145, PC3 and LNCaP | Array Express: E-MTAB-4858 | [24] |
| 27 | Su | 18 samples: 2D and 3D cultures (spheroids) derived from prostate cancer cell-line LNCaP (9 samples) and prostate normal cell-line RWPE (9 samples) | GEO: GSE30304 | No article reference found |
| 28 | Zachary | 13 samples: Organoids from primary prostate epithelial cells in mono and co-culture with stromal cells (7 | GEO: GSE115052 | [25] |

| | | | | |
|---|---|---|---|---|
| | | samples), stromal cells in co-culture (4 samples) and macrodissected tumor tissue (2 samples). | | |
| 29 | Zhang | 18 samples: Cells from benign prostatic bulk (3+3 samples), basal (3+3 samples) and luminal (3+3 samples) tissue cultured in two different media. | GEO: GSE74698 | [26] |
| 30 | Blattner | 12 samples: Mouse prostate organoids with WT (6 samples) and mutated (6 samples) gene *SPOP*. | GEO: GSE94839 | [27] |
| 31 | Aytes | 384 samples: Prostate tumor tissue from mouse models with different genetic modifications. | GEO: GSE53202 | [28] |
