## Supplementary_File_2 for "The genes controlling normal function of citrate and spermine secretion is lost in aggressive prostate cancer and prostate model systems"

**Supplementary File 2: Supplementary tables and figures**

**Integrity of initial and refined citrate – spermine (CS) gene signatures.**

**Supplementary Figure 1:** CMS for the initial CS module based on the 40 normal samples from *Bertilsson.* (Dataset ID 1). **A)** The initial CMS was not significant in the *Bertilsson* (p=0.14, lognorm test), even though the normal samples CMS (0.37) was slightly higher compared to the CMS based on cancer samples (0.34). **B)** The initial CS module (blue dots) could not be validated in prostate cancer (red circles) and normal (blue circles) samples in 11 additional datasets. The CMS was not statistically significant in any of the datasets (lognorm test, Supplementary Table 3 and 4). Dataset ID 4, 5 and 8 did not include normal samples.

A)

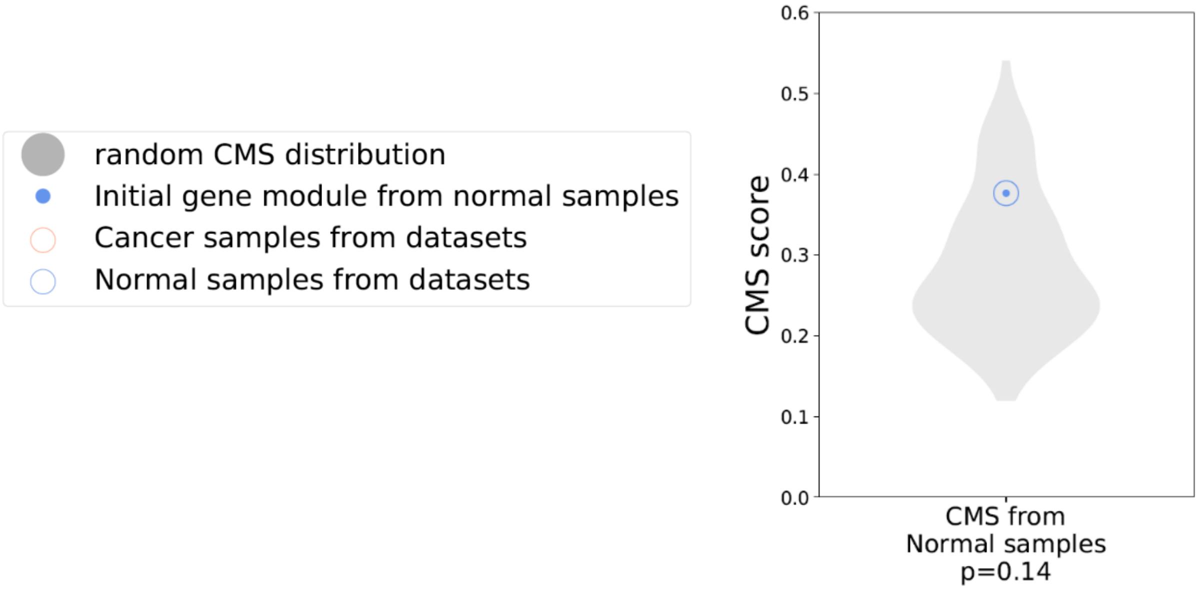


B)

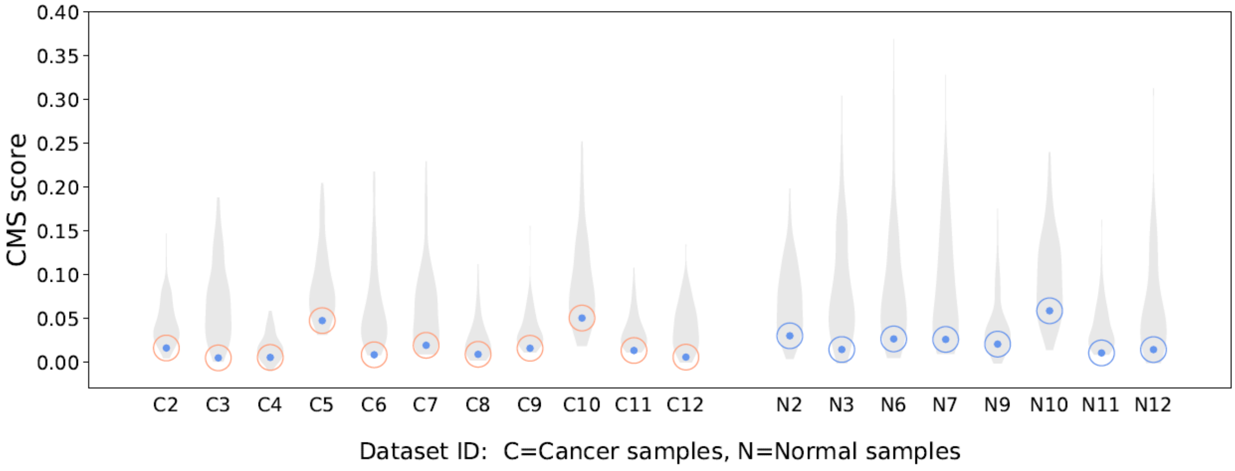

**Supplementary Table 1:** CMS for the initial **cancer** sample CS gene module from *Bertilsson* evaluated on cancer samples in 12 datasets.

| Dataset ID | Dataset Abbreviation | Cancer Samples | Number of genes | CMS score | Random average CMS score | p-value | Lognorm dist. lack-of-fit pvalue |
|---|---|---|---|---|---|---|---|
| 1 | Bertilsson | 116 | 150 | 0.35 | 0.2 | 0.008 | 0.9 |
| 2 | Chen | 65 | 115 | 0.23 | 0.05 | 0.002 | 0.4 |
| 3 | Taylor | 131 | 143 | 0.24 | 0.07 | 0.03 | 0.4 |
| 4 | Sboner | 281 | 75 | 0.13 | 0.02 | 0.02 | 0.3 |
| 5 | Erho | 545 | 142 | 0.21 | 0.1 | 0.04 | 0.3 |
| 6 | TCGA | 497 | 147 | 0.23 | 0.06 | 0.007 | 0.7 |
| 7 | CMBR | 112 | 150 | 0.21 | 0.06 | 0.02 | 0.3 |
| 8 | STCK | 94 | 150 | 0.25 | 0.04 | 0.002 | 0.4 |
| 9 | Mortensen | 36 | 143 | 0.20 | 0.04 | 0.001 | 0.3 |
| 10 | Prensner | 78 | 147 | 0.21 | 0.09 | 0.04 | 1.0 |
| 11 | Stopsack | 264 | 144 | 0.16 | 0.04 | 0.0008 | 0.9 |
| 12 | Kuner | 59 | 140 | 0.17 | 0.05 | 0.02 | 0.1 |

**Supplementary Table 2:** Supplementary Table T1: CMS for the initial **cancer** sample CS gene module from *Bertilsson* evaluated on normal samples in 9 datasets. The data from *Mortensen* are from laser dissected tissue, and the lack of significant CMS is likely due to the more homogenous epithelial tissue samples, resulting in only subtle variations in CS genes.

| Dataset ID | Dataset Abbreviation | Normal Samples | Number of genes | CMS score | Random average CMS score | p-value | Lognorm dist. lack-of-fit pvalue |
|---|---|---|---|---|---|---|---|
| 1 | Bertilsson | 40 | 150 | 0.35 | 0.09 | 0.01 | 0.9 |
| 2 | Chen | 71 | 115 | 0.25 | 0.05 | 0.01 | 0.2 |
| 3 | Taylor | 29 | 143 | 0.29 | 0.07 | 0.04 | 0.6 |
| 6 | TCGA | 52 | 147 | 0.27 | 0.05 | 0.006 | 0.3 |
| 7 | CMBR | 74 | 150 | 0.26 | 0.07 | 0.02 | 0.4 |
| 9 | Mortensen | 14 | 143 | 0.07 | 0.03 | 0.1 | 0.1 |
| 10 | Prensner | 38 | 144 | 0.23 | 0.07 | 0.02 | 1.0 |
| 11 | Stopsack | 160 | 144 | 0.17 | 0.03 | 0.0004 | 0.8 |
| 12 | Kuner | 39 | 140 | 0.22 | 0.04 | 0.004 | 0.5 |

**Supplementary Table 3:** CMS for the initial **normal** sample CS gene module from *Bertilsson* evaluated on cancer samples in 12 datasets.

| Dataset ID | Dataset Abbreviation | Cancer Samples | Number of genes | CMS score | Random average CMS score | p-value | Lognorm dist. lack-of-fit pvalue |
|---|---|---|---|---|---|---|---|
| 1 | Bertilsson | 116 | 150 | 0.08 | 0.08 | 0.4 | 0.5 |
| 2 | Chen | 65 | 86 | 0.02 | 0.04 | 0.9 | 0.8 |
| 3 | Taylor | 131 | 127 | 0.00 | 0.07 | 1.0 | 0.1 |
| 4 | Sboner | 281 | 35 | 0.01 | 0.02 | 0.8 | 0.1 |
| 5 | Erho | 545 | 128 | 0.05 | 0.09 | 0.9 | 0.8 |
| 6 | TCGA | 497 | 140 | 0.01 | 0.06 | 1.0 | 0.8 |
| 7 | CMBR | 112 | 149 | 0.02 | 0.06 | 0.9 | 0.1 |
| 8 | STCK | 94 | 149 | 0.01 | 0.03 | 0.8 | 0.6 |
| 9 | Mortensen | 36 | 132 | 0.02 | 0.04 | 0.9 | 0.8 |
| 10 | Prensner | 78 | 147 | 0.05 | 0.09 | 0.8 | 0.8 |
| 11 | Stopsack | 264 | 139 | 0.01 | 0.04 | 0.9 | 1.0 |
| 12 | Kuner | 59 | 132 | 0.01 | 0.04 | 1.0 | 0.1 |

**Supplementary Table 4:** CMS for the initial **normal** sample CS gene module from *Bertilsson* evaluated on normal samples in 9 datasets.

| Dataset ID | Dataset Abbreviation | Normal Samples | Number of genes | CMS score | Random average CMS score | p-value | Lognorm dist. lack-of-fit pvalue |
|---|---|---|---|---|---|---|---|
| 1 | Bertilsson | 40 | 150 | 0.34 | 0.27 | 0.2 | 0.4 |
| 2 | Chen | 71 | 86 | 0.03 | 0.07 | 0.9 | 0.1 |
| 3 | Taylor | 29 | 127 | 0.01 | 0.08 | 1.0 | 0.7 |
| 6 | TCGA | 52 | 140 | 0.03 | 0.09 | 0.9 | 0.9 |
| 7 | CMBR | 74 | 149 | 0.03 | 0.09 | 0.9 | 0.4 |
| 9 | Mortensen | 14 | 131 | 0.02 | 0.04 | 0.7 | 0.9 |
| 10 | Prensner | 38 | 144 | 0.06 | 0.10 | 0.8 | 0.5 |
| 11 | Stopsack | 160 | 139 | 0.01 | 0.04 | 1.0 | 0.4 |
| 12 | Kuner | 39 | 132 | 0.01 | 0.06 | 0.8 | 0.1 |

**Validation of citrate – spermine gene signature.**

**Supplementary Figure 2:** CS signature ssGSEA scores on spatial transcriptomics data from *Berglund* (dataset ID 20). The corresponding 12 pathological tissue images can be found in *Supplementary Information – Supplementary Figure 1b* in Berglund et al. (ref).

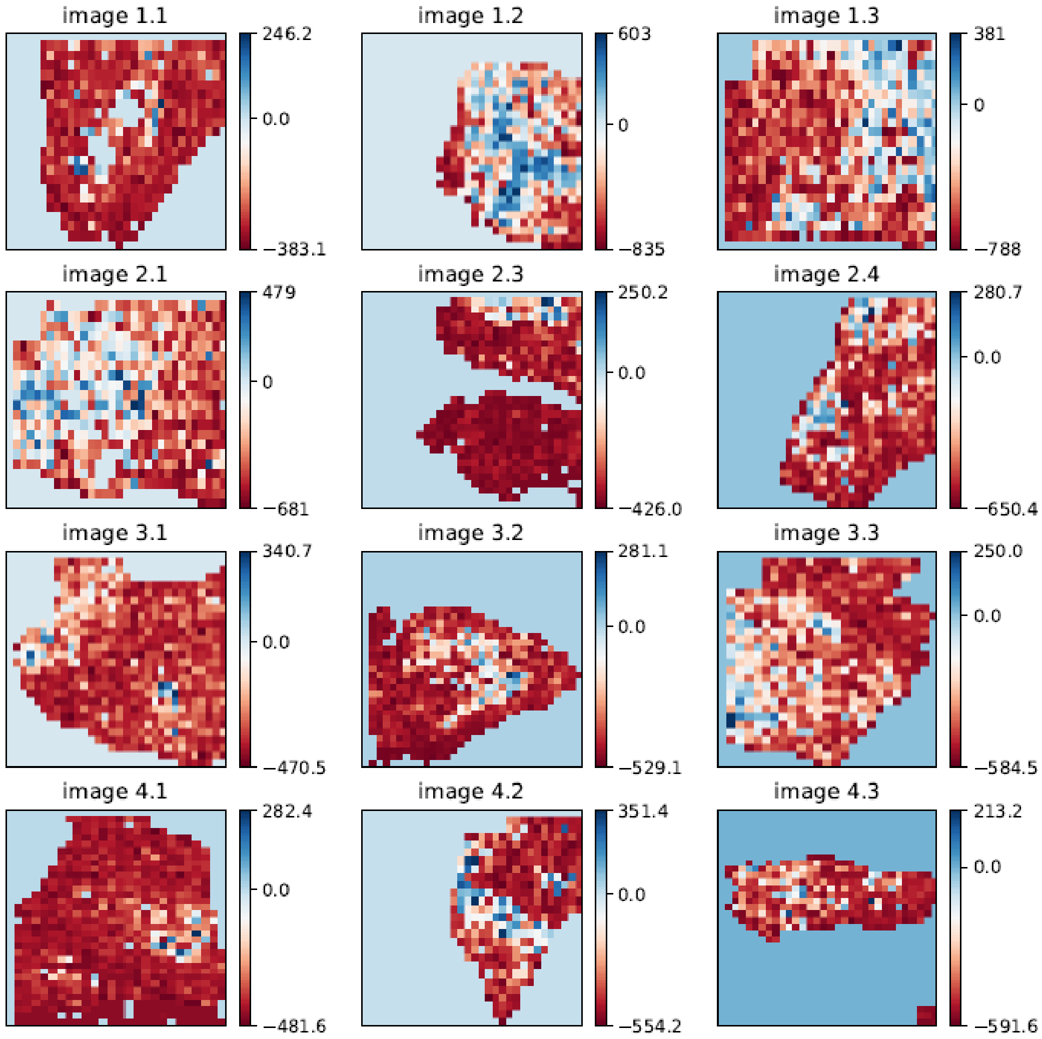

**Citrate – spermine (CS) signature, tumor grade and metastasis.**

**Supplementary Figure 3:** Stroma signature ssGSEA scores for prostate metastatis, cancer and normal prostate samples in 7 datasets. The scores were centered and normalized to range 0-1 before plotting to visualize similarities between datasets better. Normal prostate samples generally contain higher amount of stroma than cancer samples, which confounds differential expression analysis of CS signature scores involving normal prostate samples.

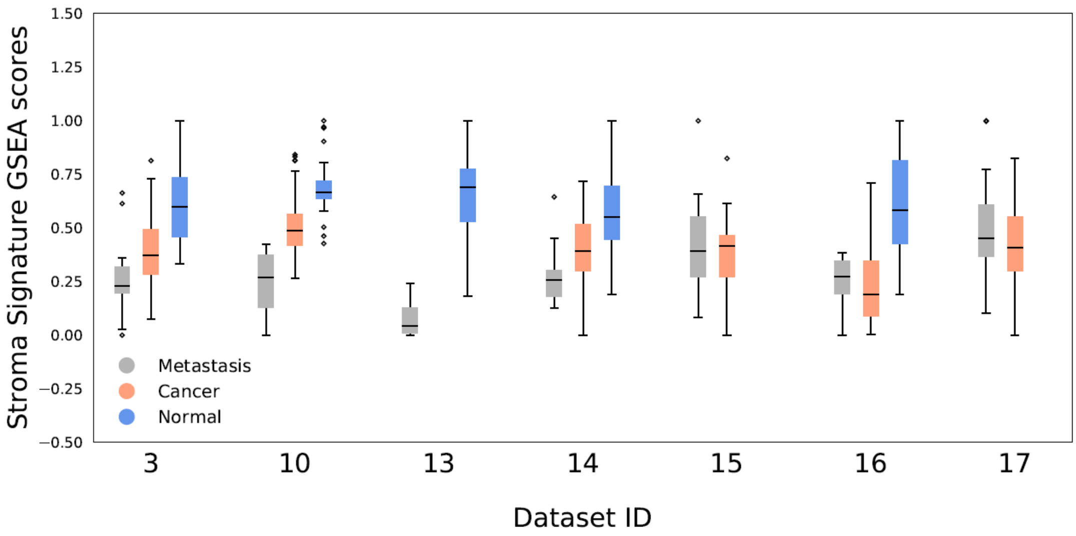

**Genes and targets**

**Supplementary Figure 4:** Significant Gene Ontology Terms from **A**) DAVID and **B**) Enrichr, for the 150 genes in CS gene signature.

**A): DAVID:**

| Category | Term | | Count | P_Value | Benjamini |
|---|---|---|---|---|---|
| **Annotation Cluster 1** | **Enrichment Score: 6.64** | | **Count** | **P_Value** | **Benjamini** |
| UP_SEQ_FEATURE | region of interest:Alpha | RT | 6 | 6.6E-9 | 2.8E-6 |
| UP_SEQ_FEATURE | region of interest:Beta | RT | 6 | 6.6E-9 | 2.8E-6 |
| UP_SEQ_FEATURE | metal ion-binding site:Divalent metal cation; cluster B | RT | 6 | 1.2E-8 | 2.5E-6 |
| UP_SEQ_FEATURE | metal ion-binding site:Divalent metal cation; cluster A | RT | 6 | 1.2E-8 | 2.5E-6 |
| INTERPRO | Metallothionein, vertebrate, metal binding site | RT | 6 | 1.9E-8 | 6.4E-6 |
| SP_PIR_KEYWORDS | metal-thiolate cluster | RT | 6 | 2.0E-8 | 5.5E-6 |
| INTERPRO | Metallothionein superfamily, eukaryotic | RT | 6 | 3.2E-8 | 5.4E-6 |
| INTERPRO | Metallothionein, vertebrate | RT | 6 | 3.2E-8 | 5.4E-6 |
| PIR_SUPERFAMILY | PIRSF002564:metallothionein | RT | 6 | 6.9E-8 | 6.1E-6 |
| SP_PIR_KEYWORDS | cadmium | RT | 5 | 2.4E-7 | 3.3E-5 |
| SP_PIR_KEYWORDS | chelation | RT | 5 | 2.4E-7 | 3.3E-5 |
| SP_PIR_KEYWORDS | acetylated amino end | RT | 9 | 1.1E-6 | 7.6E-5 |
| GOTERM_MF_FAT | cadmium ion binding | RT | 5 | 1.2E-6 | 4.6E-4 |
| SP_PIR_KEYWORDS | metal binding | RT | 5 | 1.2E-5 | 6.8E-4 |
| GOTERM_MF_FAT | copper ion binding | RT | 6 | 3.7E-4 | 2.8E-2 |
| SP_PIR_KEYWORDS | copper | RT | 5 | 9.9E-4 | 3.9E-2 |
| **Annotation Cluster 2** | **Enrichment Score: 3.45** | | **Count** | **P_Value** | **Benjamini** |
| GOTERM_MF_FAT | cofactor binding | RT | 11 | 7.7E-5 | 1.4E-2 |
| SP_PIR_KEYWORDS | nad | RT | 9 | 1.1E-4 | 5.3E-3 |
| GOTERM_MF_FAT | coenzyme binding | RT | 9 | 2.2E-4 | 2.7E-2 |
| GOTERM_MF_FAT | NAD or NADH binding | RT | 4 | 8.5E-3 | 3.7E-1 |
| **Annotation Cluster 3** | **Enrichment Score: 3.13** | | **Count** | **P_Value** | **Benjamini** |
| KEGG_PATHWAY | Valine, leucine and isoleucine degradation | RT | 8 | 5.4E-7 | 3.9E-5 |
| KEGG_PATHWAY | Fatty acid metabolism | RT | 7 | 5.3E-6 | 1.9E-4 |
| KEGG_PATHWAY | Propanoate metabolism | RT | 5 | 4.6E-4 | 1.1E-2 |
| KEGG_PATHWAY | beta-Alanine metabolism | RT | 4 | 2.0E-3 | 3.5E-2 |
| KEGG_PATHWAY | Butanoate metabolism | RT | 4 | 6.9E-3 | 7.9E-2 |
| KEGG_PATHWAY | Tryptophan metabolism | RT | 3 | 7.8E-2 | 4.1E-1 |
| KEGG_PATHWAY | Lysine degradation | RT | 3 | 9.2E-2 | 4.4E-1 |
| **Annotation Cluster 4** | **Enrichment Score: 2.99** | | **Count** | **P_Value** | **Benjamini** |
| GOTERM_BP_FAT | carboxylic acid catabolic process | RT | 8 | 3.5E-5 | 1.9E-2 |

### B): Enrichr - Pathways

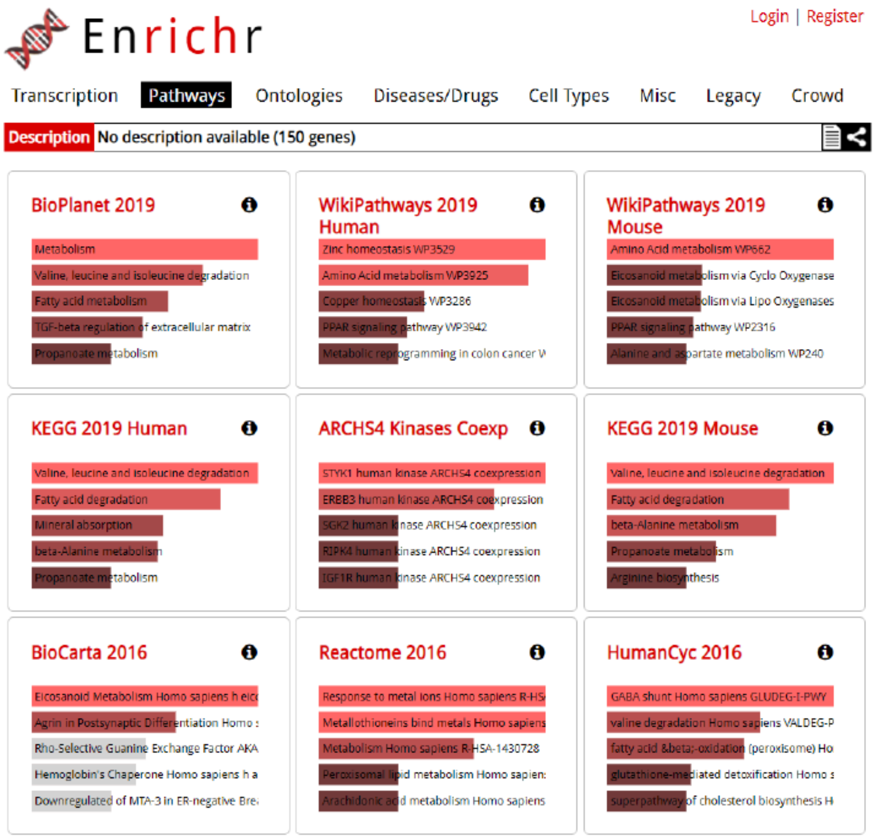


### Enrichr - Ontologies

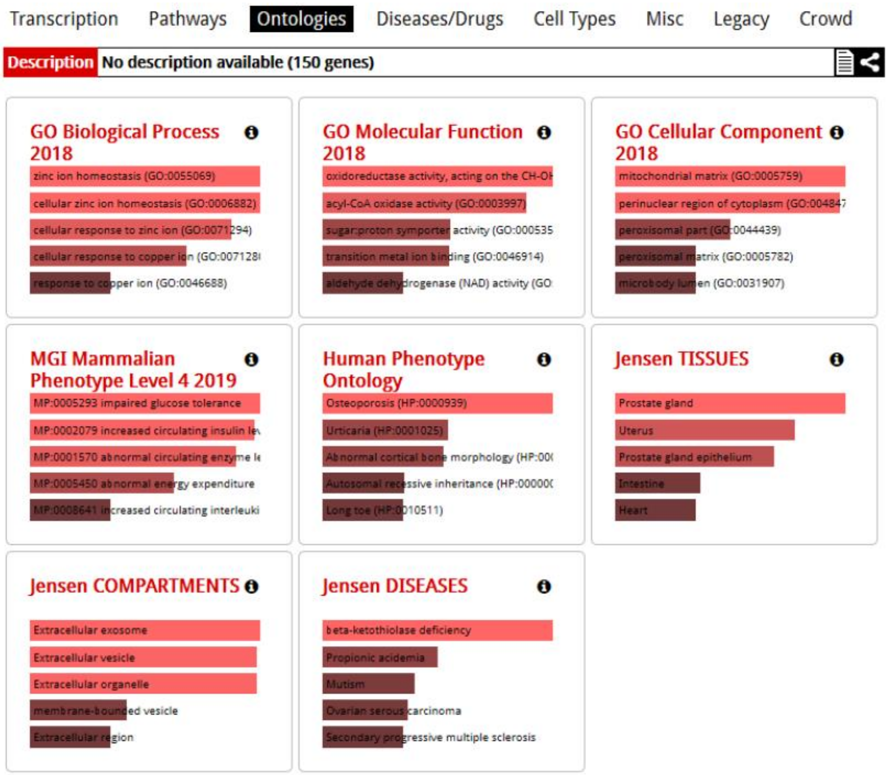

**Supplementary Figure 5:** Network analysis of the 150 genes in the CS gene signature. The network is a consensus over datasets 1-12.

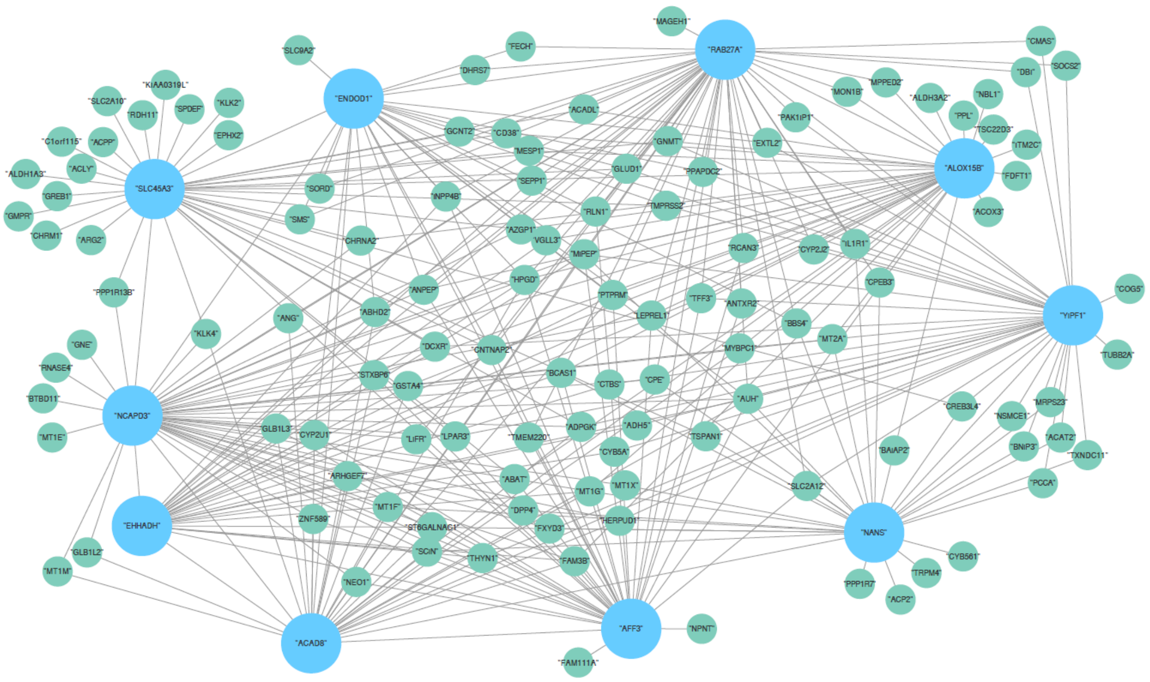

### Model Systems

**Supplementary Figure 6:** CS signature ssGSEA scores for all tissue types in cohort *Taylor* (dataset ID 3), including the four prostate cancer cell-types DU145, PC3, LNCaP and VCaP (one sample reach). The ssGSEA scores for cell-lines relative to other tissue types is similar to the *Prensner* dataset. Androgen responsive LNCaP and VCaP cell-types score higher than androgen resistant DU145 and PC3.

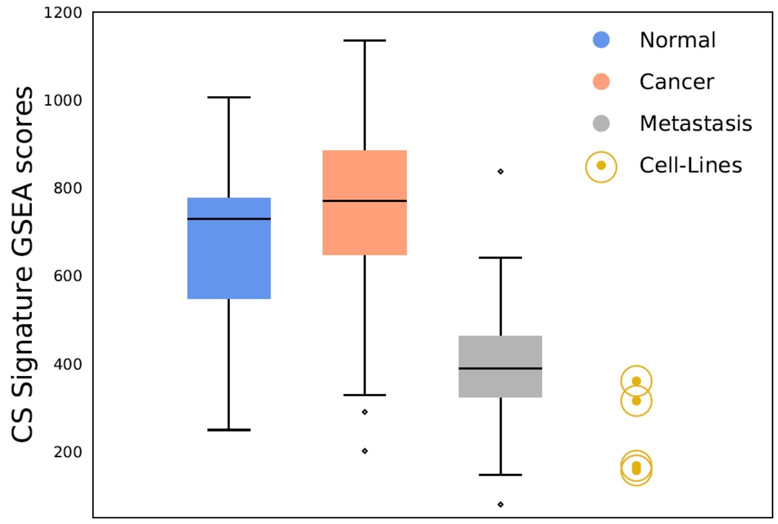

**Supplementary Figure 7:** CS Signature ssGSEA scores for the three prostate cancer cell-types LNCaP, PC3 and DU145 in *Søgaard* (dataset ID 26). There are 12 samples for each cell-type. The ssGSEA scores are highly reproducible within each cell-type, and the relative difference between the cell-types are similar to the differences observed in *Prensner* and *Taylor*, demonstrating that the assessment of CS signature enrichment of cell-lines is robust across cohorts.

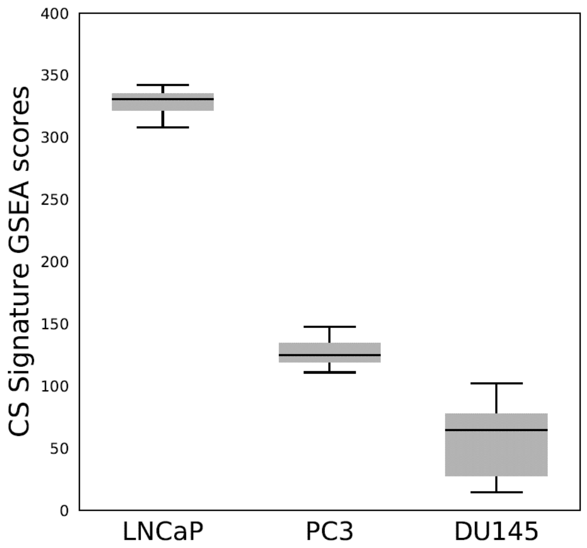


**Supplementary Figure 8:** CS signature ssGSEA scores in 622 cancer cell-lines from *E-MTAB-2706* (dataset ID 25). Androgen responsive cell-line LNCaP have higher ssGSEA score than androgen resistant cell-lines DU145 and PC3, which score similar to cell-types from other cancers. The results are similar to those observed in the *CCLE* dataset.

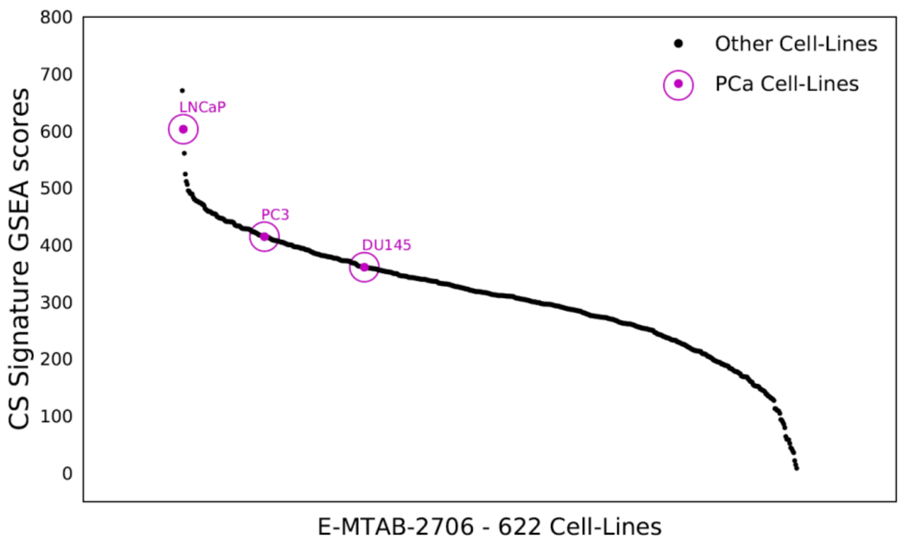

**Supplementary Figure 9:** Adapted CS signature ssGSEA scores between the *Prensner* and *Taylor* datasets. The ssGSEA scores shows that the adaptation procedure enable comparison of ssGSEA scores between different datasets. The Taylor dataset contains one sample from each of the cell-types DU145, PC3, LNCaP and VCaP.

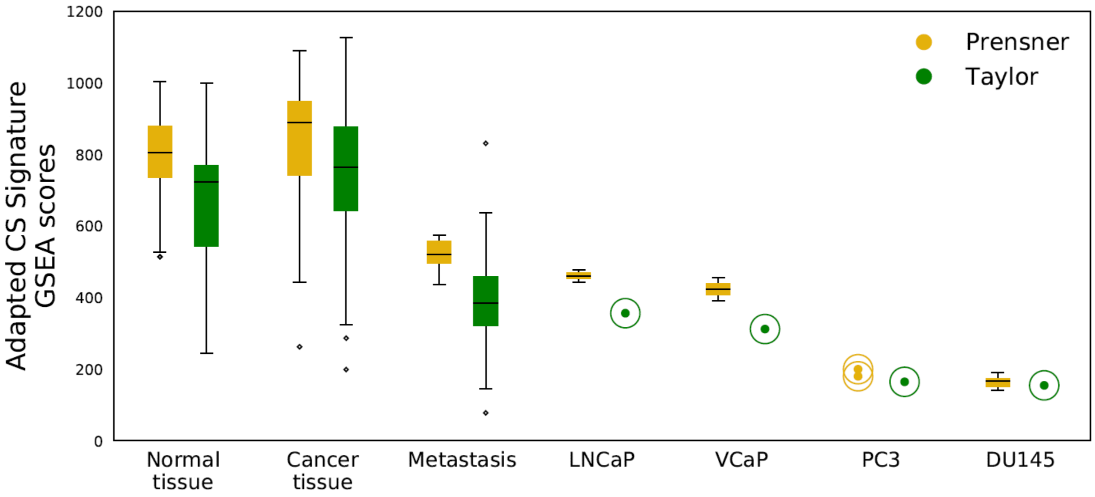


**Utility of signature refinement**

**Supplementary Figure 10:** CS signature ssGSEA scores from 96 metastatic samples (9 from prostate) in the *Hsu* dataset using the initial CS signature before refinement. Without refinement, the CS signature is not able to separate all prostate metastatic samples from other metastatic samples.

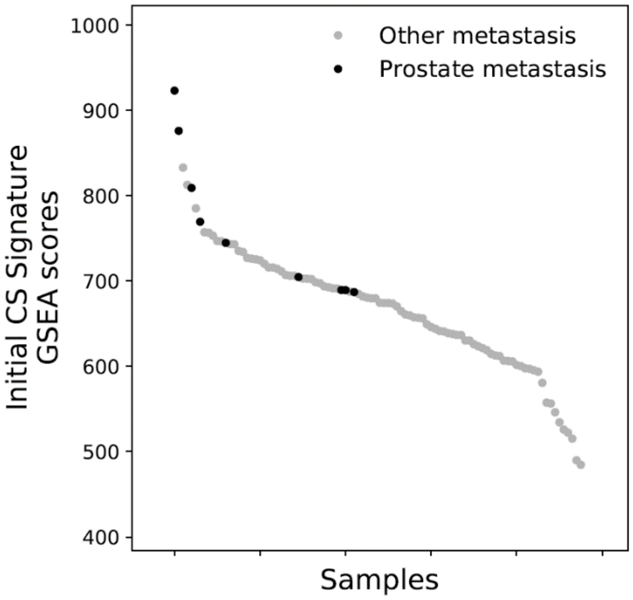

**Supplementary Figure 11:** CS signature ssGSEA scores from the *FANTOM* dataset using the initial CS signature before refinement. Without refinement, the single Prostate Adult Tissue sample do not obtain the highest ssGSEA score.

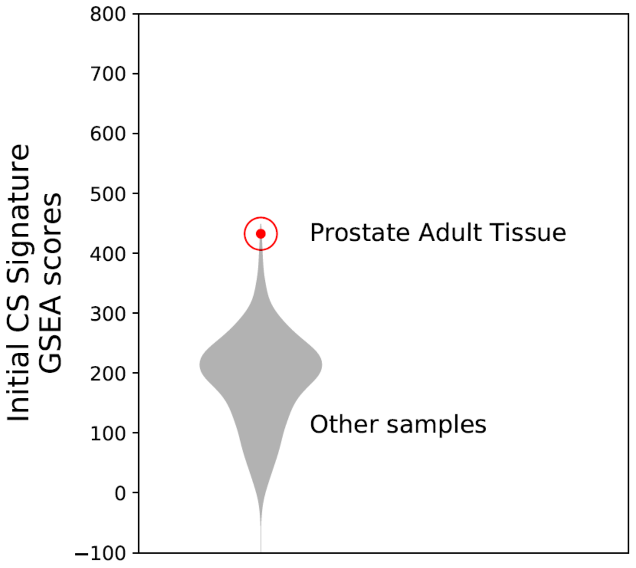


**Supplementary Figure 12:** Improvement of CS ssGSEA scores after refinement for cancer and normal samples in the *TCGA-complete* dataset. The ssGSEA score improvements are more pronounced in prostate cancer (*PRAD*) compared to other cancers. This shows that the refinement procedure for the CS signature is most effective for prostate. In addition, the results show that increased ssGSEA scores after refinement depend on the tissue type, and is not a general bias from the refinement procedure.

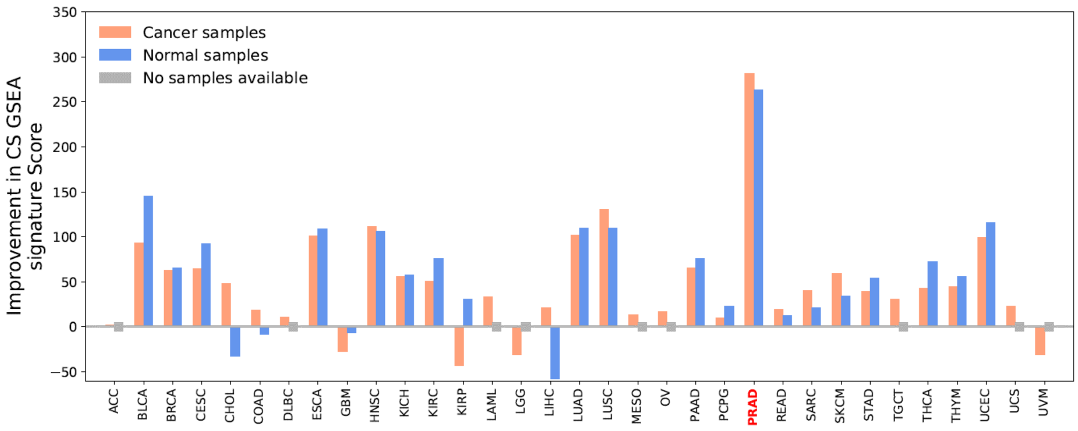

**Supplementary Figure 13:** Refined gene signature average GSEA scores for 37 metabolite and lipid-signal signatures evaluated for all 33 cancers in the *TCGA-complete* dataset. Of all metabolites and lipids, citrate and spermine have the most elevated prostate cancer GSEA scores relative to other cancers. This shows, 1) That different gene signatures with different properties are produced for different metabolites, and 2) That gene signatures for citrate are spermine are created specifically in the prostate TCGA samples, and not samples from other TCGA cancers.

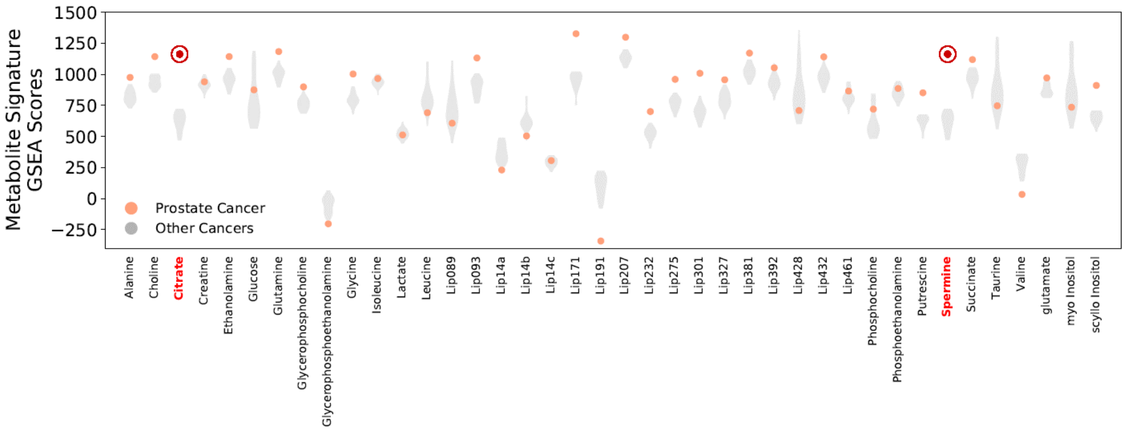


**Supplementary Figure 14:** Refined gene signature average GSEA scores for 37 metabolite and lipid-signal signatures evaluated for all 33 cancer in the *TCGA-complete* dataset. The figure is based on Supplementary Figure 13, but now the average GSEA score for other cancers have been subtracted from the prostate cancer GSEA score to emphasize the metobolite and lipid signals with the largest difference. This figure highlights whether a particular metabolite is regulated by correlated genes unique for prostate.

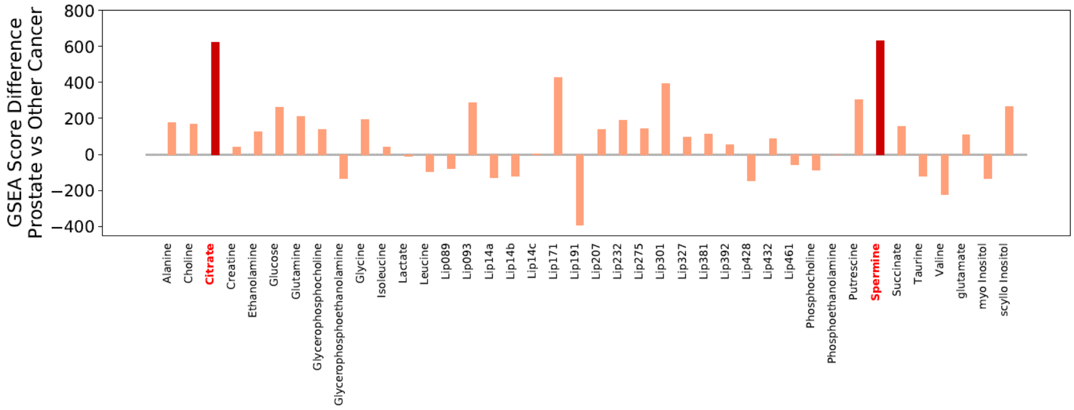


**Supplementary Figure 15:** Same as Supplementary Figure 13, but now for normal samples for 24 tissues in the *TCGA-complete* dataset. The results are the same as for the cancer samples.

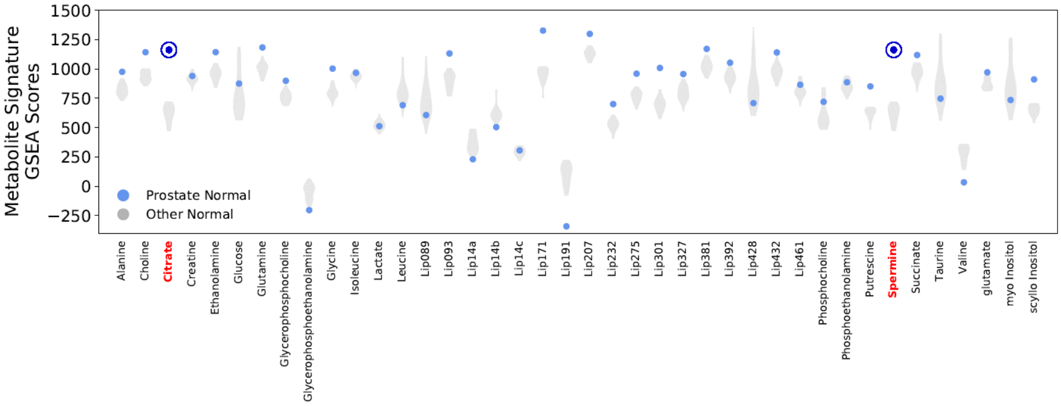

**Supplementary Figure 16:** Same as Supplementary Figure 14, but now for normal samples for 24 tissues in the *TCGA-complete* dataset. The results are the same as for the cancer samples.

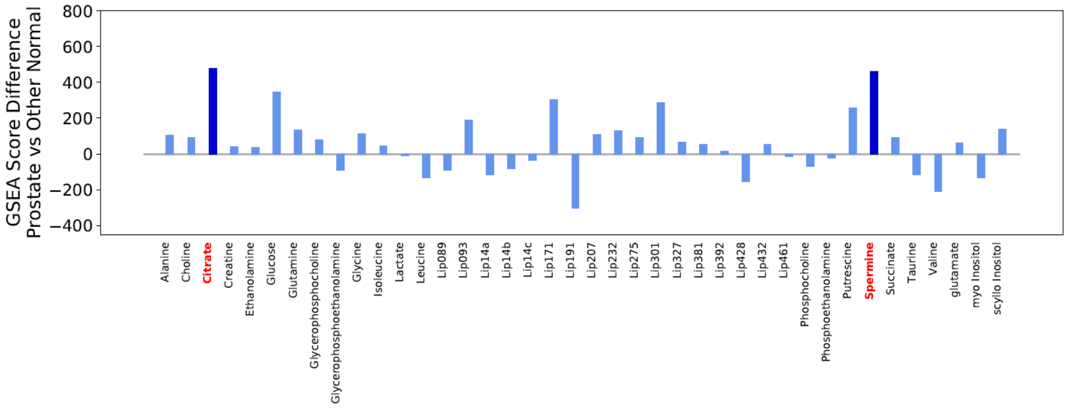


**Supplementary Figure 17:** Refined Metabolite signature GSEA score ranks for the one Prostate Adult Tissue samples in the *FANTOM* dataset (1829 samples in total) after signature refinement. Four metabolites recieved the highest rank (rank 0) for prostate adult tissue; citrate, spermine, putrescine, and the lipid signal Lip301. The ranks before signature refinement for these signals where 3, 4, 16 and 11 respectively, demonstrating that refinement improved the prostate specificity of the CS gene signature. A high rank would indicate that metabolites (or lipids) are differently regulated in prostate tissue compared to other tissues and cell-lines.

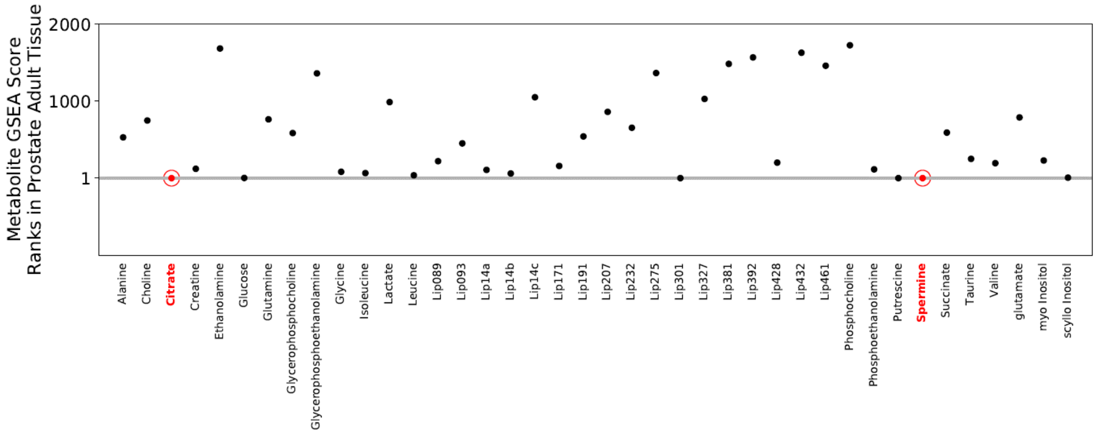


**Supplementary Figures and Tables end.**
